# An evolutionary strategy for highly efficient secretion of multidomain Staphylococcal protein A: N- to C-terminal stability gradient

**DOI:** 10.64898/2026.05.31.729144

**Authors:** Andrew Hagarman, William R. Franch, Aulane Mpouli, Bruce R. Donald, Terrence G. Oas

**Affiliations:** KBI Biopharma, Durham, NC 27704, United States; 2109 N Buchanan Ct, Arlington, VA 22207, United States; Department of Biochemistry, Duke University, Durham, NC 27710, United States; Department of Chemistry, Duke University, Durham, NC 27708, United States; Department of Computer Science, Duke University, Durham, NC 27708, United States

**Keywords:** protein folding, staphylococcal protein A, thermodynamic stability, multidomain proteins, protein secretion

## Abstract

Surface factors of *Staphylococcus aureus* have become therapeutic targets, including staphylococcal protein A (SpA), a key contributor to virulence. SpA displays five nearly identical helical (*≈*60 aa) antibody-binding domains on the bacterial surface with high affinity (*K*_*d*_ *≈* 0.4 − 4 nM). Folding free energies have been determined for only domains E and B. Here we used intrinsic fluorescence-detected urea denaturation to measure the folding thermodynamics of each domain in isolation and in the native multidomain context (SpA-N, the N-terminal half of the mature protein). Domains D, A, B, and C have identical stabilities in isolation and in SpA-N, confirming independent folding; E domain is stabilized *≈*1 kcal/mol by its multidomain context. To determine role of the conserved linker between domains D through C we used fluorescence-detected urea denaturation of 1–5 B-domain repeat constructs. Analysis of these data using a nearest-neighbor matrix model demonstrated unambiguously that adjacent domains are thermodynamically uncoupled. We also find a steep N-to C-terminal gradient of increasing domain stability (0.97 *±* 0.05 to 5.69 *±* 0.28 kcal/mol), which has not been reported for any repeat protein. We propose this gradient facilitates efficient co-translational secretion of SpA through the SecYEG translocon: the least stable E domain enters the channel first, lowering the energetic barrier to translocation of this highly abundant surface protein.

**Significance:** Staphylococcal protein A (SpA) is a highly expressed virulence factor of *Staphylococcus aureus* that evades host immunity by binding antibodies with high affinity. SpA displays on the bacterial surface five nearly identical small helical domains that are connected by highly flexible linkers. We measured the folding thermodynamics of each domain in isolation and in their native multidomain context. We unambiguously demonstrated that adjacent domains are thermodynamically uncoupled. We also discovered a steep N-to C-terminal gradient of increasing domain stability. We propose this gradient facilitates efficient post-translational secretion of SpA through the SecYEG translocon: the least stable domains enter first, lowering the energetic barrier to translocation for the most highly expressed protein in *S. aureus*.

## Introduction

Staphylococcus aureus is responsible for illnesses ranging from non-serious to life threatening. This bacterium has a myriad of surface factors that contribute to its virulence(1, 2) including the antibody binding protein, staphylococcal protein A (SpA)(3). Recent efforts in therapeutic treatment of *S. aureus* associated illness have focused on protein A(4). Fig. 1 shows a domain representation of SpA. The C-terminal half is responsible for sorting and cell wall attachment and contains the following segments: a variable repeating octapeptide region(5); an LPXTG attachment motif(6); a hydrophobic region; and a positively charged C-terminal tail(7). The other half of SpA contains five antibody binding domains(8) and an N-terminal signal sequence(9) used in the secretion process. SpA is secreted post-translationally and cleaved by sortase (SrtA) between the T-G peptide bond in the LPXTG attachment motif and anchored to the cell wall(10) so that on the exterior of the bacterium the five N-terminal antibody binding domains are displayed. These domains bind the Fc region of antibodies with high affinity (*K ≈* 0.4 − 4 nM)(11), which inhibits opsonization and phagocytosis(12).

**Fig 1.**
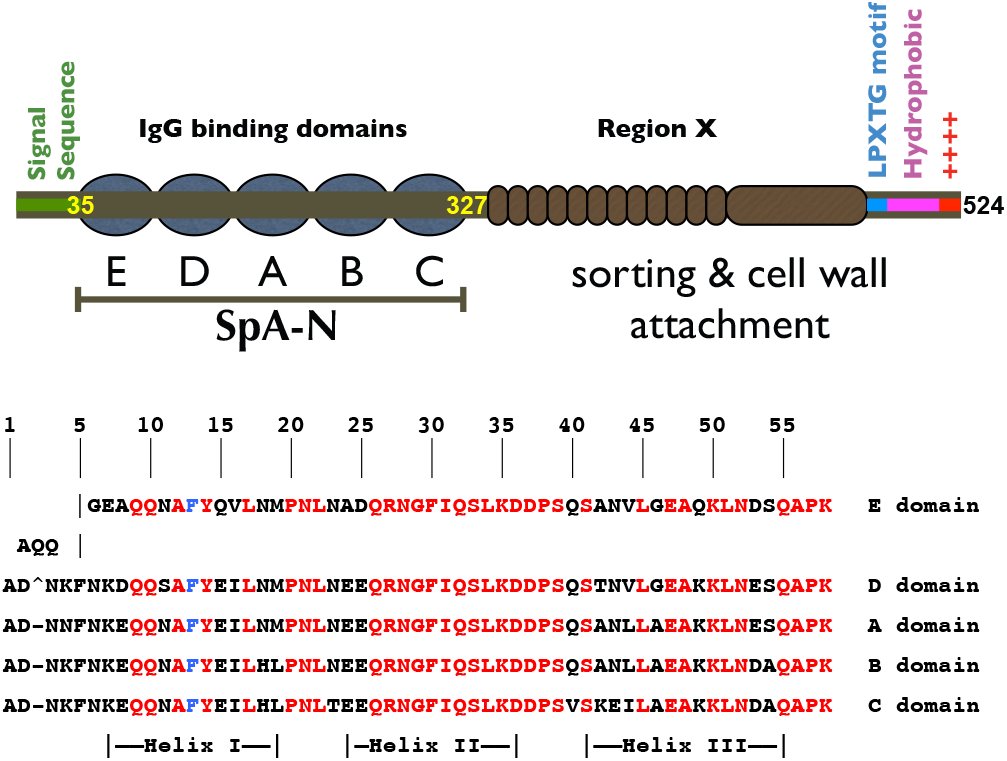
Domain representation of protein A showing the N-terminal signal sequence, the five antibody binding domains (E, D, A, B, C), region X (the presumably disordered variable repeat octapeptide region), the LPXTG cell wall attachment motif, a hydrophobic sequence, and the C-terminal positively charged tail. The primary amino acid sequence of SpA-N is listed with each domain aligned vertically. Residues conserved among all domains are highlighted in red. The site of tryptophan substitutions (F13, etc.) is colored in blue.

For the studies described here, we used the residues of SpA corresponding to its N-terminal half (without the signal sequence), referred to hereafter as SpA-N. SpA-N contains the five antibody binding domains designated E, D, A, B, and C(8). The individual domains are three helix bundles that are nearly identical in sequence, as seen in Fig. 1 and Table 1. Helix I consists of 8-12 residues and has 6 identical residues across all five domains. Helix II consists of 12-13 residues with the last 11 residues being identical in all five domains. The first two residues in Helix II differ only in the E domain. Helix III, which Sato, et al. determined using phi analysis to be the most stable(13), yet the least formed in the transition state, consists of 15 residues and is the least conserved of the helices. Only seven of the residues in Helix III are identical in all five domains. The turns connecting HI-HII and HII-HIII are both four residues long. Both turn regions have nearly identical sequences across all five domains. The isolated B domain of protein A (BdpA) has been the focus of considerable previous work including the following topics: thermodynamics and kinetics of folding(13–20); antibody binding(21, 22); phi-analysis(13, 20); solution NMR structure(16, 22–24); and crystal structures(21, 22, 25, 26). BdpA is a popular molecule for simulations of fast protein folding(27–31) and has also been used as a system to test and calibrate force fields in molecular dynamics simulations because it exhibits a two-state folding mechanism and has a simple topology(32). In contrast, there have been few studies of the other four domains in isolation or their native multidomain context(33, 34).

**Table 1.** Pairwise SpA-N domain sequence identity in percent.

| Domain | EdpA | DdpA | AdpA | BdpA | CdpA |
| --- | --- | --- | --- | --- | --- |
| EdpA | - | 83 | 83 | 79 | 72 |
| DdpA | 83 | - | 91 | 84 | 79 |
| AdpA | 83 | 91 | - | 91 | 83 |
| BdpA | 79 | 84 | 91 | - | 91 |
| CdpA | 72 | 79 | 83 | 91 | - |

Multidomain proteins comprise a significant portion of proteomes and represent more than two-thirds of eukaryotic proteins(35). Linker regions usually connect the domains, vary in length and structure and can contribute to function(36, 37). Domains connected by long, flexible linkers are likely to act independently of their neighbor. This sort of independence is necessary for domains to perform different functions. Conversely, domains connected by short, structured linkers are likely to produce a cooperative effect on their neighboring domains. Two coupled domains have interfacial interactions that contribute to the folding free energy of each domain. These energetically linked interfacial residues do not necessarily have to be located in the linker to be coupled to the folding of an adjacent domain.

It is intriguing to consider the role of thermodynamic inter-domain coupling in the stability of multidomain proteins. Thermodynamic coupling means that the folding free energy of a domain is directly affected by that of its neighbor. This effect can be large and in some cases related to function. For example, in the tandem R16–R17 construct (R1617) from chicken brain *α*-spectrin, a single Glu-to-Pro substitution corresponding to a pathogenic mutation in human spectrin (37) destabilizes the R16 domain by *≈*1 kcal/mol (35). Because the mutation lies in the helix connecting R16 and R17, it also effectively abolishes the nearly 3 kcal/mol stabilizing interaction between the two domains. Thus, this single mutation destabilizes R17 by *≈*5 kcal/mol.

Thermodynamically uncoupled domains, such as those in the I-band of titin, act essentially independently of their neighbors. Although these domains are in proximity, their linkers are stiff and neighboring domains do not interact. Scott, et al. described the I-band of titin as the “sum of its parts”(38). This may be advantageous for the I-band of titin, which is responsible for passive elasticity in muscle. Under tension, only a few Ig domains unfold thereby elongating the protein(39). The observed independence confers the ability of some domains to remain folded when their neighbors unfold. Upon releasing tension, the unfolded domains are allowed to then freely fold and escape any misfolded structures involving neighboring unfolded chains.

Other examples of multidomain thermodynamic coupling are found in the following linear repeat sequences: notch ankyrin repeats (ANK)(40, 41); tetratricopeptide repeats (TPR)(42); leucine rich repeats (LRR)(43); and hexapeptide repeats(44). These ubiquitous motifs are characterized by their incorporation in proteins as successive repeats, which are nearly identical in structure. ANK repeats and TPR encompass two alpha helices connected by turns in each subunit. LRR are typically mixed *α*/*β* structures. Hexapeptide repeats are composed of *β*-strands that form a so-called *β*-helix. Biophysical studies of ANK repeat, TPR, and LRR proteins have determined the thermodynamics of their folding. In some cases, consensus sequence repeats have been used to simplify the analysis. Both ANK and TPR consensus sequence repeats increased stability upon addition of flanking domains, indicating strong thermodynamic coupling between neighboring domains. The unfavorable intrinsic stabilities of the monomer units of each repeat are counterbalanced by a more favorable interfacial energy. Thus, the folding of the repeat proteins becomes favorable, despite the instability of the individual units. Kloss and Barrick systematically investigated leucine rich repeats from YopM by deleting C-terminal domains(43). By making deletions of C-terminal repeats they found that some repeats had little to no effect on the stability while others caused the stability to drop markedly. In this regard, deletions of the C-terminal *β*-cap, the 11th, 6th, and 5th repeats had the largest effect. These results led the authors to conclude that two regions (repeats 12-15 and 7-10) are unstable and require interfacial interactions with the C-terminal and internal (repeats 6 and 11) caps in order to fold. These cases of thermodynamic coupling underscore the importance of interfacial interactions and protein stabilization as a consequence.

Although there have been many biophysical studies of SpA domains (BdpA in particular), some disagreement exists in the literature about whether adjacent domains interact. Starovasnik, et al. reported thermodynamic results on E domain and the two-domain E–D fragment(33). The authors’ differential scanning calorimetric results show that the E domain is fairly unstable, with a *T*_*m*_ of 43 °C. The authors reported a broad transition for the E–D fragment, which is a consequence of the difference in the stabilities of each domain. Their analysis indicated that the D domain is slightly more stable than the E domain and that the two domains do not interact.

Two studies have focused on coupling of adjacent B domains. BdpA has the most identical residues compared with the other four domains (Table 1) and can be considered a consensus sequence so that repeat B domains are analogous to the aforementioned consensus repeat proteins. In the first study, Karimi, et al. compared circular dichroism detected thermal melts of single (BdpA) and double B (BBdpA) domains and concluded that there was an apparent difference in cooperativity due to the interaction of adjacent B domains(16). The authors also reported interdomain nOe’s and correlation times calculated from fluorescence depolarization measurements, which they attributed to an interaction between linked domains. More recently, Arora, et al. contradicted the results of Karimi et al. and concluded that adjacent B domains do not interact(19). Arora, et al. compared fraction denatured plots calculated from circular dichroism detected, denaturant induced unfolding of BdpA and BBdpA. The calculated fraction denatured plots for each protein perfectly overlay each other, indicating that neither the free energy of folding nor the *m*_*eq*_ value for the B domain were perturbed by adding an adjacent identical domain. However, one interpretation of this result is that the interface is disrupted at low denaturant concentrations and is thereby absent in the observable portion of the denaturation curve. In addition, the possibility exists that small interaction energies could not be detected with only one interface, in the case of two B domains, but could become more apparent by adding other domains.

In this paper we used the consensus sequence approach and extended the model to multi (1-5) B domain constructs and monitored their fluorescence as a function of urea denaturation. We globally fitted the experimental data using the homopolymer matrix method(41) and unambiguously show that adjacent B domains do not thermodynamically interact. We also measured the stabilities of the antibody binding domains in SpA-N and compared them to the stabilities of the isolated single domains to show that, except for E domain, there is no significant effect of stability on neighboring domains and that these domains do, in fact, fold independently. We also found a large and monotonic increase in domain stability, from N-to C-terminus that we propose may play a role in the efficient secretion of protein A.

## Results

### B domain repeat proteins have no interdomain coupling

#### Fitting nonlinear unfolded fluorescence baselines

As a fluorescent probe of SpA domain folding, we used a conserved F13W (BdpA numbering) substitution in each domain. This substitution has been used previously in BdpA(17, 18, 20). The F13 side chain is situated on the surface of the protein. Thus, the substituted Trp sidechain fluorophore is quenched upon unfolding. The wavelength of maximum emission increased as a function of increasing denaturant. This bathochromic shift is typical for the exposure of fluorophores to aqueous solvent.

The linear extrapolation method(45) assumes that the folding free energy, and the folded and unfolded signals are linearly dependent on denaturant concentration. Thus, there are six parameters that determine the slopes and intercepts of these lines. Often, when these parameters are simultaneously fitted using nonlinear least squares methods the covariance between the baseline parameters and the thermodynamic parameters is high.

A complication in the application of the linear extrapolation method to denaturation curves detected by intrinsic tryptophan fluorescence (275-280 nm excitation) is the non-linearity of tryptophan fluorescence vs. [Den]. As seen in Fig. 2, the fluorescence of N-acetyl-tryptophanamide (NATA) clearly deviates from linearity at high urea concentrations. NATA is a viable model for tryptophan in an unfolded protein although changes in quantum yield compared to tryptophan in an unfolded protein could concomitantly change the curvature of the baseline. Nevertheless, we find that NATA works perfectly in this case, which is exemplified by the fits in Fig. 2 and the agreement between the two green and blue curves shown. We fitted the NATA titration curve to a sum of linear and exponential components as seen in Eq. 1.

**Fig 2.**
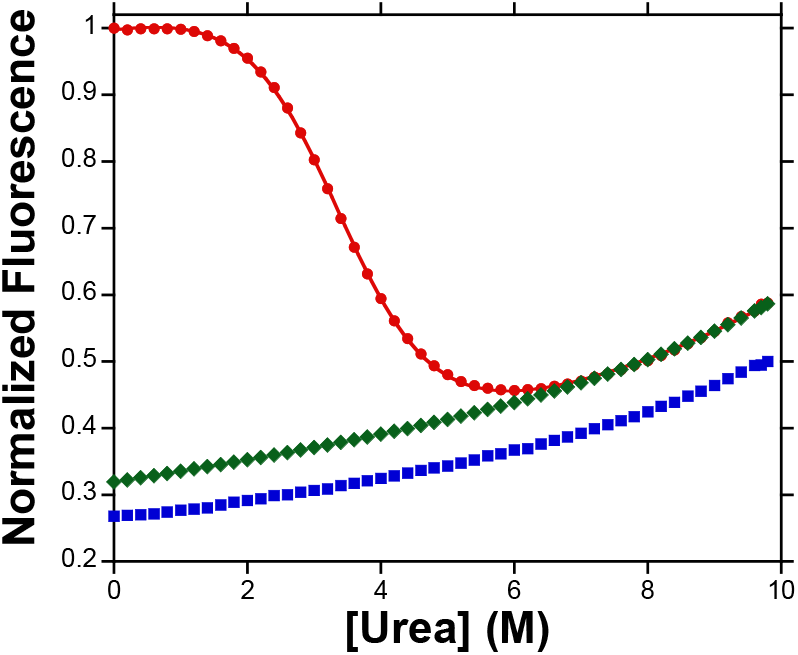
Fluorescence detected urea denaturation curve of DdpA (red circles), which highlights the nonlinear unfolded baseline. The denaturation curves were normalized to the signal in 0 M urea as explained in the text. The titration of N-acetyl tryptophanamide (NATA) (blue squares) was used to obtain the parameters for the nonlinear contribution to the unfolded baseline. The fit to the DdpA denaturation curve (red line) was generated using the unfolded baseline parameters from fitting the NATA titration. The unfolded baseline (green diamonds) was generated from the fit to the DdpA denaturation curve, with A and B from Eq. 1 fixed to 0.0075 and 0.3, respectively, while Θ_U,0_ was allowed to float.

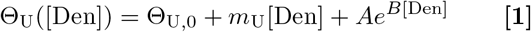

where Θ_U_([Den]) is the signal for the unfolded baseline, Θ_U,0_ is the signal of the unfolded baseline at zero denaturant, *m*_*U*_ is the slope of the linear part of the baseline, the exponential prefactor, A, accounts for the amplitude of the deviation from linearity at high urea concentrations, and B is the growth constant. To fit the unfolded base curves of SpA-N domains, we used the A (0.0075) and B (0.3) factors obtained from the fit to the NATA titration as fixed parameters that describe the non-linearity, as depicted for DdpA in Fig. 2. In contrast, fitting the unfolded DdpA base curve to a line rather than Eq. 1 gave a *m*_eq_ value of 0.77 ± 0.02 kcal/mol M-1, which is 7% lower than the best fit value (Table 2) obtained using Eq. 1. On the other hand, when we used a similar approach for the folded baseline, there was no improvement in the quality of the fit and no significant difference in the best-fit thermodynamic parameter values. It is imperative to properly model the folded and unfolded base curves while fitting denaturation data. This is especially the case when the urea dependence of the folded or unfolded signals is not well determined because of low or high stability (e.g., EdpA and CdpA). Normalizing the signal and fitting the unfolded base curve to Eq. 1 with fixed A and B values still allowed for variation in the slope of the folded baseline and the vertical position and linear slope of an unfolded base curve, which accommodated variations in instrument sensitivity and protein concentration. This fitting approach gives statistically well-determined thermodynamic parameters with low covariance.

**Table 2.** Transition midpoint (*C*_½_), free energy of unfolding in the absence of denaturant, and the linear urea dependence of the free energy of unfolding (*m*_eq_) at 25 °C for each of the isolated single domains from SpA-N. The uncertainties are 95% confidence intervals based on reproducibility analysis (see Error Analysis).

| Single Domain | E | D | A | B | C |
| --- | --- | --- | --- | --- | --- |
| $C_{1/2}$ (M) | $0.04 \pm 0.88$ | $3.39 \pm 0.22$ | $5.51 \pm 0.36$ | $6.53 \pm 0.43$ | $8.09 \pm 0.53$ |
| $\Delta G(0)$ (kcal mol<sup>-1</sup>) | $0.02 \pm 0.46$ | $2.67 \pm 0.17$ | $4.20 \pm 0.27$ | $4.79 \pm 0.31$ | $5.07 \pm 0.33$ |
| $m_{eq}$ (kcal mol<sup>-1</sup> M<sup>-1</sup>) | $-0.53 \pm 0.03$ | $-0.79 \pm 0.05$ | $-0.76 \pm 0.05$ | $-0.73 \pm 0.05$ | $-0.63 \pm 0.04$ |

#### Denaturation of 1-5 B domain repeat proteins

Thermodynamic analysis of linear repeat proteins is amenable to nearest-neighbor models that minimize the number of energy terms. Models of these proteins that contain identical consensus sequence repeats simplify the analysis further. The translational symmetry in linear repeat proteins reduces the number of energy terms necessary to describe the folding of the entire molecule to a single intrinsic domain stability and single domain-domain interaction free energy. We chose the B-domain sequence as a consensus sequence because it is most similar to the other four domains of SpA-N (Table 1.)

The B-domain repeat proteins we constructed were linked in the same way that domains A, B, and C are linked. In order to unambiguously address whether coupling exists between adjacent B-domains, we globally fit the fluorescence detected urea denaturation curves of single, double, triple, quadruple, and quintuple B domain constructs (1–5B), as shown in Figure 3, with the homopolymer matrix method(41). Briefly, the homopolymer matrix method is a matrix-based derivation of the partition function for linear repeat proteins that accounts for all possible 2^*N*^ species assuming only two thermodynamic states for each of the N domains. The key parameters are intrinsic folding free energy (*G*_folded_ − *G*_unfolded_ = Δ*G*_*i*_) of domain i and the free energy of interaction (*G*_interface_ − *G*_no interface_ = Δ*G* _+1_) between neighboring domains *i* and *i ±* 1, used here. The matrix form of the partition function for *N* domains, *Q*(*N*), is written as shown in Eq. 2.

**Fig 3.**
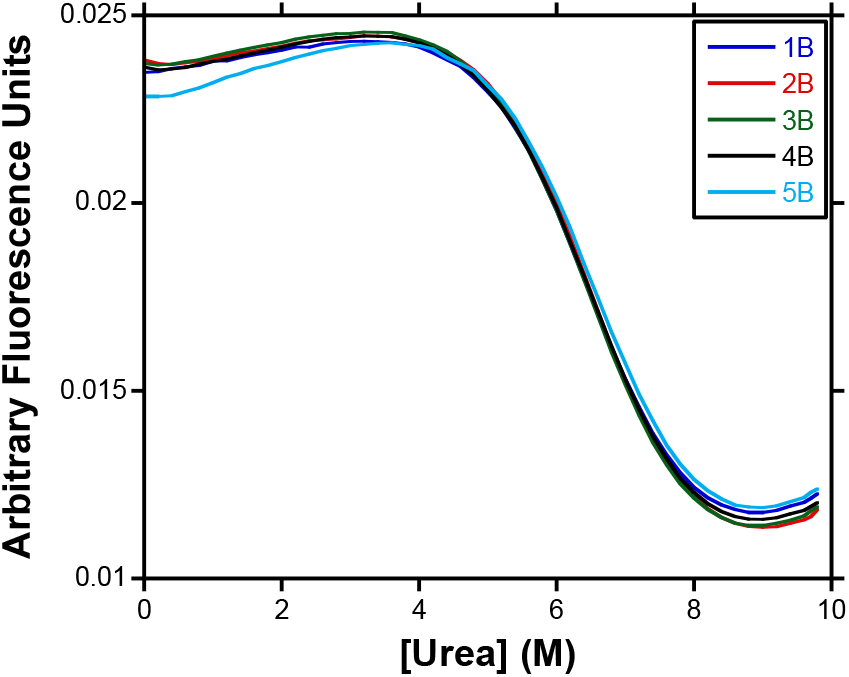
Fluorescence detected urea denaturation curves at 25 °C for single (red line), double (blue line), triple (green line), quadruple (black line), and quintuple (cyan line) B domain. The denaturation curves were normalized as explained in Materials and Methods.

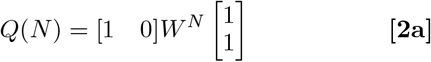

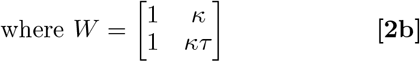

and where *W* is the statistical weight matrix, 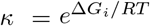, and 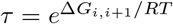. The so-called *m*_*eq*_ value (*d*Δ*G*_*i*_*/d*[Den]) is proportional to the change in solvent accessibility upon folding. Taking the product of all *W* matrices is greatly simplified by treating it as an eigenvalue problem, where *W*^*N*^ = *TD*^*N*^ *T* ^−1^. *D* is a diagonal matrix of eigenvalues and *T* is an invertible matrix of its eigenvectors. After calculating the partition function, we fit the experimental denaturation curves in terms of fractional folded repeats, *θ*, to Eq. 3.

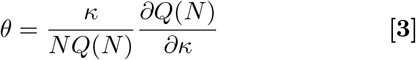

It is qualitatively apparent that there is no coupling between adjacent B-domains because all five denaturation curves overlay on each other with small differences in the folded and unfolded baselines due to experimental uncertainty. Also, the mean residue ellipticity did not significantly change by adding additional B domains, as seen in the far-UV CD spectra for all five multi-B constructs (Fig. 4). The variation in intensity of these spectra followed no discernable trend and is therefore likely due to experimental uncertainty.

**Fig 4.**
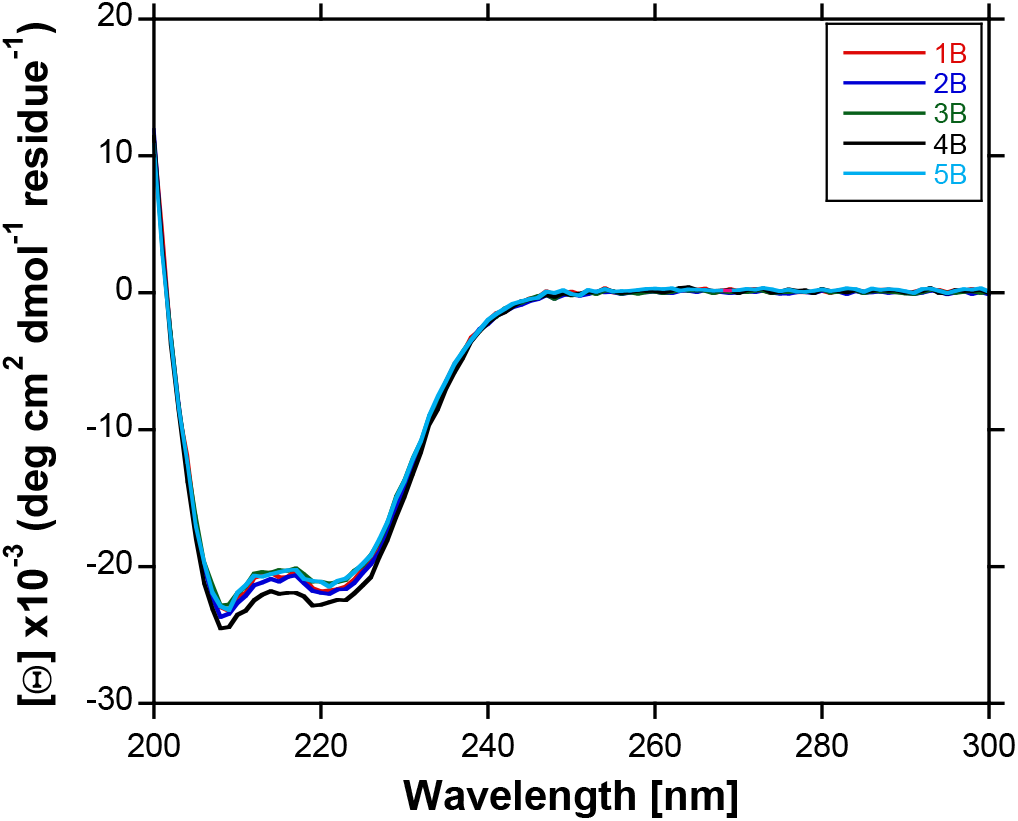
Far-UV circular dichroism spectra for single (blue line), double (red line), triple (green line), quadruple (black line), and quintuple (cyan line) B domain. The spectra are plotted in mean residue ellipticity from 200-300 nm at 25 °C.

Like the folding free energy, the coupling free energy should have a linear dependence on denaturant concentration, i.e., 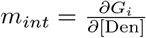. Because initial fits yielded a small coupling energy, this derivative was hard to measure experimentally. Thus, *m*_*int*_ was held fixed at 0 to obtain a best-fit coupling free energy in the absence of denaturant.

Global fits of all five denaturation curves with the homopolymer matrix method(41) yielded a folding Δ*G*(0) of −4.63 *±* 0.13 kcal mol^−1^ and a *m*_*eq*_ value of 0.71 *±* 0.02 kcal mol^−1^ M^−1^. The intrinsic stability of Δ*G*_*i,i*+1_ is −0.03 *±* 0.14 kcal mol^−1^, which consequently confirms the original conclusion of Arora et al. that adjacent B-domains are not thermodynamically coupled(19). In contrast, Karimi, et al. reported interdomain nOe’s for their BB protein and attributed this observation to interdomain interaction(16). While this may be the case, it does not necessarily indicate thermodynamic coupling between domains. The spatial requirements for nOe’s are almost certainly different than that of an interface with thermodynamic coupling. Constraints for the latter are unclear. Also, there are differences in the sequence of BB protein used by Karimi, et al. (referred to in their paper as FB2)(16) and the one used here and in Arora, et al.(19) The Karimi, et al. sequence added an additional Met residue at the N-terminus of the first domain and six extra residues (ADNKGS) at the C-terminus of the second domain(16). These residues could have broken the symmetry of the two domains and affected the circular dichroism detected temperature melt of FB2 compared with the single domain protein. There may also have been significant errors in the actual temperature of the samples used by Karimi, et al. because sample temperature gradients during automated thermal denaturation can lead to large errors at elevated temperatures, in this case, near the transition temperature. To resolve these discrepancies we measured the circular dichroism detected thermal denaturation of B and BB, in a 1.0 cm cuvette with proper stirring and a calibrated temperature sensor, seen in Supporting Information (Fig. S1). The thermal denaturation curves overlay well and indicate no change in cooperativity of the unfolding of BB, although the unfolded baselines are poorly defined due to the high denaturation temperature. It is possible that the differences observed by Karimi, et al. only exist at very elevated temperatures where the driving forces for domain-domain interactions may be substantially different. We believe that our chemical denaturation data are closer to physiological conditions and therefore more biologically relevant.

The lack of thermodynamic coupling we observed is consistent with our small angle X-ray scattering (SAXS) studies, which showed that the scattering curves of multiple B-domain constructs 2B–5B could be well-fit to an excluded volume pearl necklace model, which depicted the linker between domains as flexible gaussian chains(46). This conclusion is consistent with the order parameters we measured by ^15^N-^1^H NMR relaxation experiments, which indicated high order parameters for the core domains and low order parameters for the 6 linker residues (KADNKF)(47). Both these structural studies indicate a lack of structural coupling between domains, which supports our conclusion that there is no thermodynamic coupling between domains.

#### The domains in SpA-N have increasing stabilities from the N-to C-terminus

The SpA antibody binding domains have similar sequences, as shown in Table 1. The pairwise sequence identities range from 72% identical (E and C) to 91% identical (D and A, A and B, B and C). The structures of the individual domains are similar also, based on similarity among the published structures (22, 23, 34) and the far-UV CD spectra of each domain (Fig. 5). The CD spectra of domains D, A, B and C are identical within experimental error. The EdpA CD spectrum is less intense than the other domains because at room temperature it is rather unstable and approximately 10-15% unfolded. Interestingly, despite the differences in stability and CD spectra, the solution structure of the E domain(34) is nearly superimposable with the structures of domains D(25) and B(22, 23).

**Fig 5.**
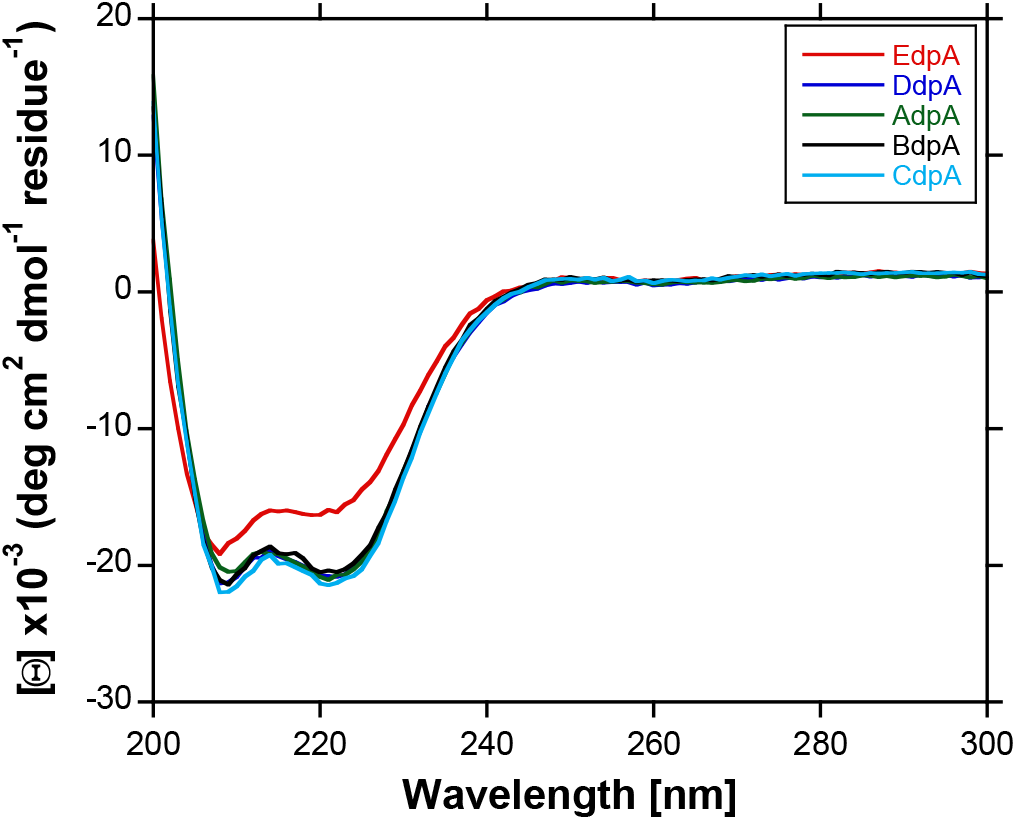
Far-UV circular dichroism spectra for EdpA (blue line), DdpA (red line), AdpA (green line), BdpA (black line), and CdpA (cyan line). The spectra are plotted in mean residue ellipticity from 200-300 nm at 25 °C.

Despite this high degree of sequence and structural similarity, the stabilities of these five domains vary over a wide range. We obtained the fluorescence urea denaturation curves of each isolated domain with tryptophan substituted at the position corresponding to residue 13 (see Fig. 1) in each domain. This substitution is mildly destabilizing for BdpA (0.25 kcal/mol)(19). The near identity of the flanking sequence in the other domains suggests that the same holds for them as well. Thus, the results observed with the tryptophan labeled proteins are likely to be very similar to the wild type sequences. Fig. 6A shows the raw data and fits for the five isolated domains from SpA-N. Fig. 6B shows the respective fraction denatured curves. We used a two-state thermodynamic model and the fitting procedure explained above to generate the fits.

**Fig 6.**
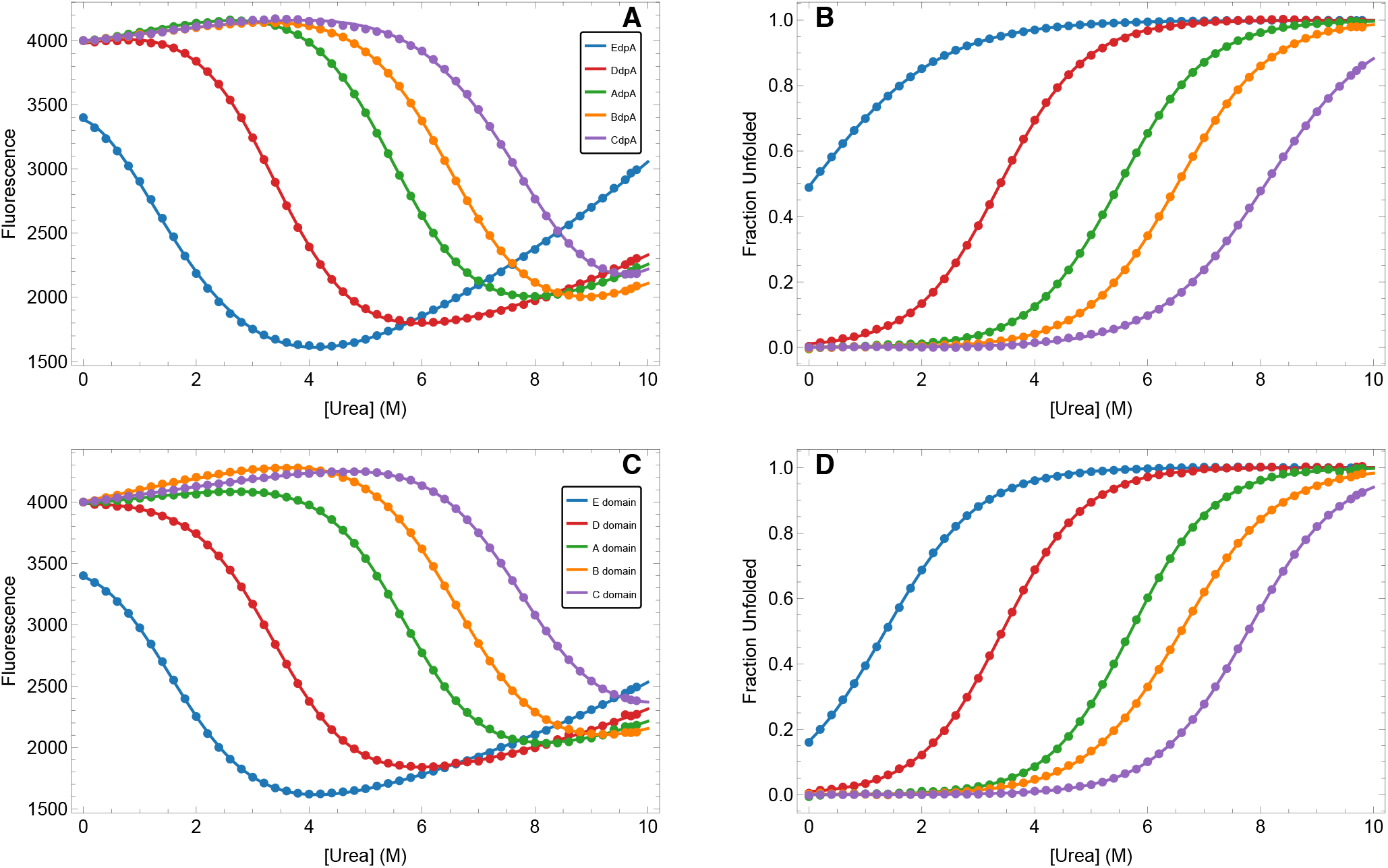
The fluorescence-detected denaturation curve of each isolated domain is nearly identical to that of the same domain in full-length SpA-N. Colors correspond to domain: E (blue), D (red), A (green), B (orange), C (purple). (A) Fluorescence detected urea denaturation curves at 25 °C (points) and fits (solid lines) for isolated domains. (B) Fraction denatured plots for isolated domains. (C) Fluorescence detected urea denaturation curves at 25 °C (points) and fits (solid lines) for domains in SpA-N. (D) Fraction denatured plots for domains in SpA-N.

Our results show that the stabilities of these domains span a wide range (see Table 2.) The E domain of protein A (EdpA) is unstable. Its equilibrium stability as an isolated domain is −0.02 *±* 0.46 kcal/mol at 25 °C. Fig. 1 shows that EdpA is 5 residues shorter than the other domains because the first three residues after the signal peptidase cleavage site do not conform to the consensus sequence of the other four domains and are not likely to be structured. The D domain (DdpA) is more stable than E, in agreement with the results from Starovasnik, et al. (33) with a stability of -2.67 ± 0.13 kcal/mol. The A domain (AdpA) and D domain have 91% sequence identity, yet AdpA is considerably more stable at -4.20 ± 0.21 kcal/mol. This increasing stability trend continues, where the B and C domains have stabilities of -4.79 ± 0.24 kcal/mol and -5.07 ± 0.25 kcal/mol, respectively. Thus, there is a gradient of increasing stability from the N to C-terminus, in the same order as the sequence. This stability gradient is apparent in Fig. 7. To the best of our knowledge, this type of stability gradient in a repeat protein has not been reported before and may be an evolved trait. It is interesting to consider its possible biological utility, as discussed below.

**Fig 7.**
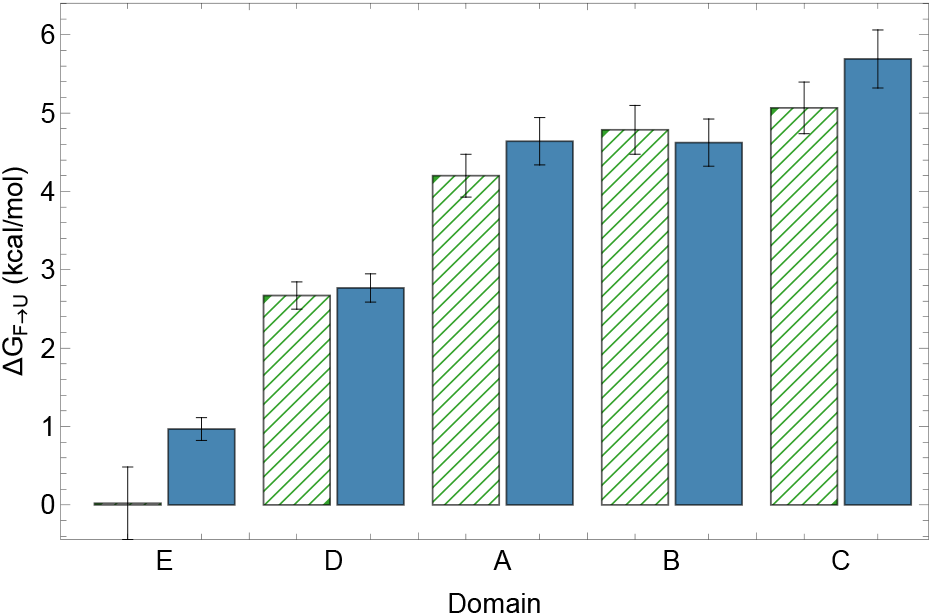
All domains except E have the same stability within error as isolated domains and as part of full-length SpA-N. Unfolding free energies at 25 °C in the absence of denaturant for isolated single domains are shown as green striped bars on the left side for each domain and SpA-N domains are shown as blue solid bars on the right side for each domain. Domains are indicated on the X-axis. The error bars are 95% confidence intervals based on reproducibility analysis (see Error Analysis).

#### Except for E domain, the stability of each isolated domain is similar to that of the same domain in full length SpA-N

To determine the stability of each domain within intact SpA-N, we substituted the phenylalanine at position 13 in a given domain with a tryptophan and left the remaining four domains unchanged (see Fig. 1.) This approach yielded five different SpA-N variants with domain specific probes that differ only by which domain contained the F13W substitution. The tryptophan fluorescence as a function of denaturant concentration gives a denaturation curve for the tryptophan labeled domain in its SpA-N context. We chose 295 nm excitation so that the fluorescence spectra and denaturation curves do not contain significant contributions from tyrosine fluorescence in the other four domains. The raw data and fits are shown in Fig. 6C. The corresponding fraction denatured curves are shown in Fig. 6D. The lack of coupling in the folding of the nB proteins suggests that there is also no coupling in SpA-N. If this were the case, then fits to the data using a two-state thermodynamic model would allow a comparison of the stability of each domain in its isolated and SpA-N contexts. As stated by Batey et al. and references within, if the thermodynamic parameters obtained from each context agree, then the lack of coupling is verified(35). The folding free energies extrapolated to zero denaturant for the domains in SpA-N are shown in Table 3, along with *m*_eq_-values and their estimated errors. The denaturation curves and fits are shown in Fig. 6.

**Table 3.** Transition midpoint (*C*_½_), free energy of unfolding in the absence of denaturant, and the linear urea dependence of the free energy of unfolding (*m*_eq_) at 25 °C for each of the domains in their multidomain context. The uncertainties are 95% confidence intervals based on reproducibility analysis (see Error Analysis).

| SpA-N Domain | E | D | A | B | C |
| --- | --- | --- | --- | --- | --- |
| $C_{1/2}$ (M) | $1.35 \pm 0.18$ | $3.43 \pm 0.22$ | $5.70 \pm 0.37$ | $6.59 \pm 0.43$ | $7.78 \pm 0.51$ |
| $\Delta G(0)$ (kcal mol<sup>-1</sup>) | $0.97 \pm 0.15$ | $2.77 \pm 0.18$ | $4.64 \pm 0.30$ | $4.62 \pm 0.30$ | $5.69 \pm 0.37$ |
| $m_{eq}$ (kcal mol<sup>-1</sup> M<sup>-1</sup>) | $-0.72 \pm 0.05$ | $-0.81 \pm 0.05$ | $-0.81 \pm 0.05$ | $-0.70 \pm 0.05$ | $-0.73 \pm 0.05$ |

By comparing the values reported in Tables 2 and 3, it is apparent that for domains D, A, B and C, each isolated domain stability is within the estimated error for the same domain in SpA-N. This comparison can be seen graphically in Fig. 7, which supports the conclusion that these domains in SpA-N fold independently with no detectable interaction free energy. In addition, the stability gradient observed in the isolated domains also holds in the SpA-N context. The only domain in SpA-N whose stability differs from that of the isolated domain is E. Its presence in SpA-N apparently stabilizes it by *≈* 1 kcal/mol. This stabilization may be the consequence of the long linker between E and D, which is KADAQQNKF, 3 residues longer than the other linkers (see Discussion).

#### The stabilities of domains A, B & C are insensitive to low pH

It has been appreciated for over 50 years that staphylococcal protein A is tolerant of acidification to pH values as low as 2.5 and this property has been exploited to elute antibodies from SpA resins(48). Our experiments to measure the thermodynamic stability of SpA domains, A, B and C provide an explanation for this acid tolerance. Fig. 8 shows the free energy of denaturation for each of these domains at 4 acidic pH values. This observation may also be biomedically significant because it has been established that *Staphylococcus aureus* can survive the low pH environment of phagolysosomes(49) as one of the consequences of its acid tolerance(50).

**Fig 8.**
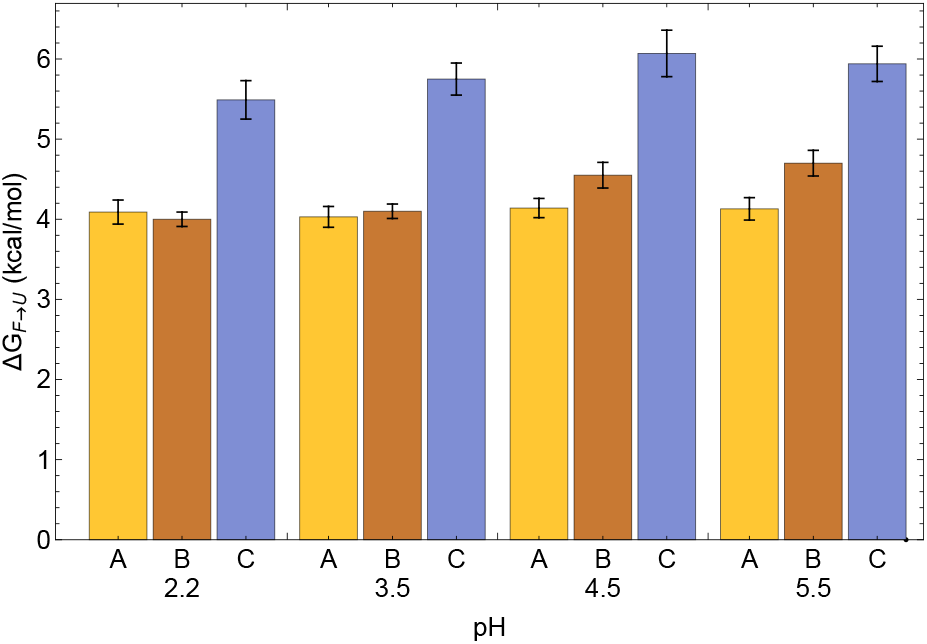
The free energy of denaturation of domains A, B and C at various acidic pH values. Domain A is indicated in burnt orange, B in brown and C in blue. Error bars represent 95% confidence levels obtained from the fits of urea denaturation curves.

#### Denaturation surfaces of each domain provide detailed thermodynamics of folding equilibria

The CD denaturation surfaces and a discussion of the fits of each of the five isolated domains are shown in the supplementary information and Fig.s S4-S8. The fitted thermodynamic parameter estimates and the parameter estimates derived from them at 25 °C (*i*.*e*., Δ*H*, Δ*S, T* Δ*S*, Δ*G* and *m*_*eq*_), are listed in Table 4. Although many of these estimates are quite uncertain, particularly for EdpA, the estimates of Δ*G* and *m*_*eq*_ agree well with the values obtained above from fluorescence denaturations. The parameter estimates also provide useful insights about the thermodynamic driving forces for the folding of these small three helix bundle proteins. The favorable free energy of folding at 25 °C is due to a favorable enthalpy of folding that is greater than the unfavorable entropy. The temperature of maximum stability (*T*_*hs*_) and the concentration of urea needed to reach Δ*G* = 0 (*C*_*hs*_) both increase with domain stability, indicating that the *G*(*T*, [Den]) curve shifts both to the right and up as stability increases.

**Table 4.** Best fit thermodynamicun parameter estimates for unfolding at 25 °C from individual domain CD denaturation surfaces (see Supporting Information Figures S4-S8).

| Parameter | units | E | D | A | B | C |
| --- | --- | --- | --- | --- | --- | --- |
| $T_{hs}$ | °C | $0. \pm 8.$ | $-1.4 \pm 1.5$ | $4.2 \pm 0.7$ | $5.3 \pm 1.5$ | $13.3 \pm 1.5$ |
| $C_{hs}$ | M | $1.6 \pm 0.5$ | $4.09 \pm 0.04$ | $6.25 \pm 0.04$ | $6.96 \pm 0.06$ | $7.79 \pm 0.11$ |
| $\Delta C_p$ | kcal mol<sup>-1</sup> K<sup>-1</sup> | $0.81 \pm 0.24$ | $0.51 \pm 0.04$ | $0.617 \pm 0.025$ | $0.59 \pm 0.05$ | $0.53 \pm 0.06$ |
| $\Delta H$ | kcal mol<sup>-1</sup> | $17. \pm 10.$ | $23.1 \pm 1.7$ | $16.7 \pm 1.2$ | $20.8 \pm 2.2$ | $21.2 \pm 2.5$ |
| $\Delta S$ | cal mol<sup>-1</sup> K<sup>-1</sup> | $55. \pm 37.$ | $66. \pm 6.$ | $43. \pm 4.$ | $52. \pm 7.$ | $53. \pm 8.$ |
| $T\Delta S$ | kcal mol<sup>-1</sup> | $16. \pm 11.$ | $19.8 \pm 1.7$ | $12.7 \pm 1.2$ | $15.5 \pm 2.1$ | $15.7 \pm 2.3$ |
| $\Delta G$ | kcal mol<sup>-1</sup> | $0. \pm 12.$ | $3.3 \pm 1.9$ | $4.0 \pm 1.5$ | $5.3 \pm 2.7$ | $5.5 \pm 3.2$ |
| $m_{eq}$ | kcal mol<sup>-1</sup> M<sup>-1</sup> | $-1. \pm 4.$ | $-1.0 \pm 0.4$ | $-0.72 \pm 0.22$ | $-0.82 \pm 0.34$ | $-0.7 \pm 0.4$ |
| $h$ | kcal mol<sup>-1</sup> M<sup>-1</sup> | $1.6 \pm 3.0$ | $-2.33 \pm 0.27$ | $-0.63 \pm 0.16$ | $-1.32 \pm 0.25$ | $-1.93 \pm 0.29$ |
| $s$ | cal mol<sup>-1</sup> K<sup>-1</sup> M<sup>-1</sup> | $9. \pm 10.$ | $-4.6 \pm 0.9$ | $0.3 \pm 0.5$ | $-1.7 \pm 0.8$ | $-4.1 \pm 0.9$ |

#### SpA-N denaturation curves are broad and non-sigmoidal

To aggregate our results for the individual domains, we have both thermally and chemically denatured full-length SpA-N. As expected, the resulting denaturation curves are broad, relatively featureless, and non-sigmoidal. To compare the predicted and observed denaturation curves, we have used the parameters in Tables 4 and 3 to predict the aggregate fraction unfolded curves and used those predictions in combination with linear baseline models to fit the folded and unfolded baselines (see Materials and Methods). Using these baselines, we converted the signal in the denaturation curves to fraction unfolded, and the result is shown as data points in Figure 9. The agreement between the fraction unfolded curves coming from the individual domain parameters and the observed denaturation curves is striking. The unfolding of SpA-N comprises many partially folded microstates. On the scale of the entire protein, unfolding is non-cooperative. However, we have previously demonstrated that each domain unfolds in a two-state cooperative way (17, 19, 51).

**Fig 9.**
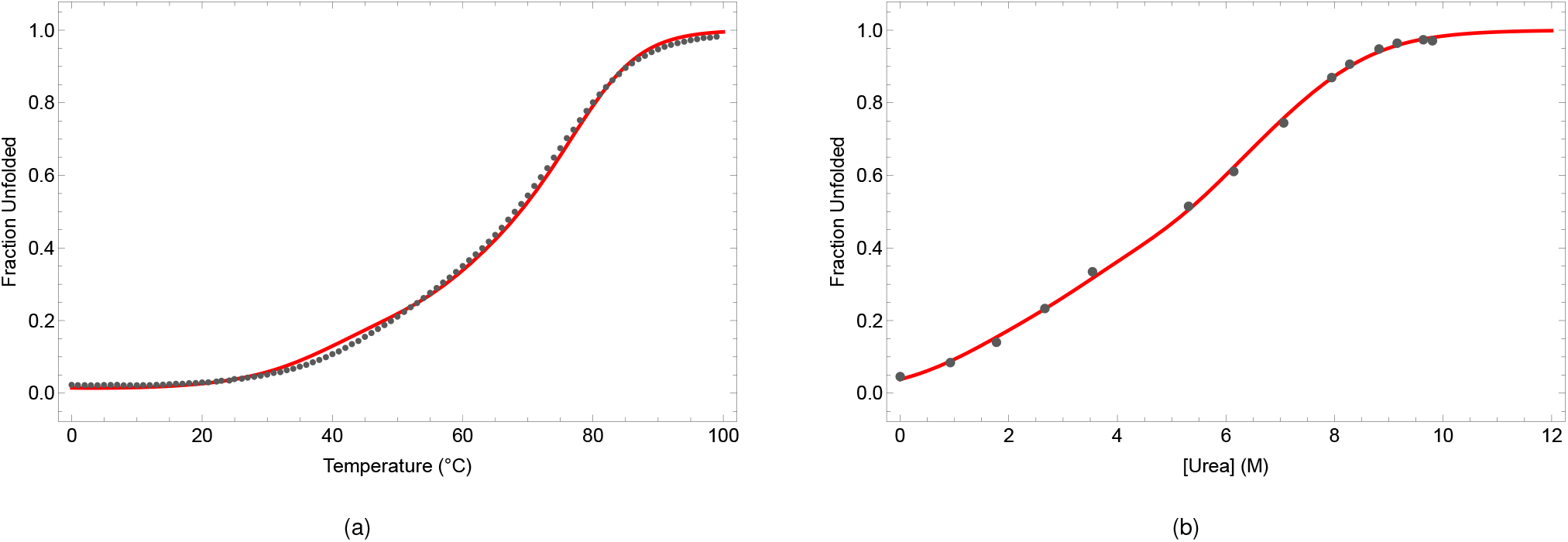
Predicted and observed CD denaturation curves for SpA-N. a) The raw CD thermal denaturation data were fit to linear folded and unfolded baselines using the model parameters listed in Table 4 (see Figure S9) to give the observed fraction denatured curves (blue data points.) The red curve shows the calculated aggregate fraction of SpA-N denatured based on the model parameters in Table 4. b) The raw CD urea denaturation data were fit to linear folded and unfolded baselines using the model parameters listed in Table 3 (see Figure S10) to give the observed fraction denatured curves (blue data points.) The red curve shows the calculated aggregate fraction of SpA-N denatured based on the model parameters in Table 3.

The results discussed above establish that each domain folds independently, which allows a straightforward statistical thermodynamic model to predict the fraction unfolded curves for SpA-N. Using the parameters in Tables 4 and 3, it is possible to compute the equilibrium constants for unfolding at any given temperature or urea concentration, respectively. The partition function for computing the populations of each of the 32 possible microstates can be written as follows:

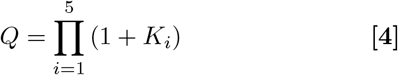

where *K*_*i*_ represents the temperature- and urea-dependent equilibrium constant for unfolding of the *i*th domain.

The variable *i* represents the index of the domain in the sequence EDABC. There are 32 terms in this equation when multiplied out. Each term in this partition function represents the statistical weight of a microstate, and the population of each microstate can be calculated by dividing its statistical weight by *Q*.

The microstate label is the 5-character string FFFFF, where F is replaced by U in every position that has a corresponding *K*_*i*_ in its statistical weight. For example, microstate FFFFF has the term equal to 1, and UUUFF has the term equal to *K*_E_*K*_D_*K*_A_. The sum of the population of each microstate multiplied by one-fifth of the number of its U domains gives the fraction unfolded depicted in Figure 9. In addition, it is possible to compute the population of each microstate as a function of temperature or urea.

Figures S11 and S12 depict these populations as a function of temperature and urea, respectively, and demonstrate that the most populated microstate progresses as FFFFF → UFFFF → UUFFF → UUUFF → UUUUF → UUUUU while sample conditions go from low temperature/urea to high. The population of each successive microstate reaches a maximum at progressively higher temperatures or urea, thus giving rise to the broad denaturation curves observed. Of the remaining 26 non-sequentially unfolded microstates, only 4 in thermal denaturation and 2 in urea denaturation exceed 5% at any condition. Overall, the populated microstates roughly follow the stability gradient observed from the N-to C-terminal domains. This distribution of partially folded microstates is consistent with the hypothesis for secretion discussed below.

## Discussion

A majority of evolved proteins comprise more than one domain (52). Thus, understanding the process by which such proteins achieve and maintain the folded structure of these domains is crucial to understanding protein folding in general. As mentioned above, the domains of some multidomain proteins are coupled thermodynamically suggesting that their folding process is itself coupled (43). However, many examples of multiple independent folding domain (MIFD)(19) proteins have been characterized (19, 38, 52). There has been much recent discussion of the evolutionary origins of multidomain folding and the role that translation plays in the process(53, 54). Here, we have described a bacterial surface protein with 4 of 5 domains that fold independently, presumably during and after translation. The B domain has been shown to fold in less than a millisecond (17, 29, 51). Thus, the independent folding of these domains in protein A almost certainly occurs as soon as they are synthesized and before the protein has been released from the ribosome. Because unfolding is also fast, many unfolding/refolding cycles must occur before the newly synthesized protein reaches the SecYEG apparatus for translocation across the *S. aureus* plasma membrane(55). It should be noted that *S. aureus* has no secB gene, which in Gram-negative bacteria acts as a post-translational chaperone to keep newly synthesized proteins unfolded prior to translocation. Thus, SpA arrives at the translocon already folded but needs to be unfolded prior to translocation. Catipovic, et al. demonstrated that the ATP-consuming unfoldase SecA uses a powerstroke unfolding mechanism and that unfolding of the translocated protein is rate limiting(56). SecA facilitates SpA secretion(55), but the role of the low stability of E domain in secretion efficiency has not been determined.

Staphylococcal protein A is the most highly expressed protein in *S. aureus*(57). For this reason, the synthesis and secretion of SpA must be highly efficient. We hypothesize that some of this efficiency is achieved through the stability gradient of the N-terminal domains of the protein. The first domain to enter the translocon after the signal sequence would be E domain. The experiments described above demonstrate that E domain is only *≈*50% folded at physiological temperatures. This property should greatly facilitate the speed at which the SpA chain is engaged with SecYEG, facilitating whatever unfolding activity SecA provides. Considering the large abundance on the surface of *S. aureus* cells, their rapid growth rate, and the limited availability of SecYEG translocons, the low stability of E domain is likely to be an evolutionary adaptation. By the same logic, the next domain (D) also has low stability albeit not as low as E. We hypothesize that once E and D are through the translocon, the transmembrane potential or other mechanical interactions assist in the unfolding of the more stable A, B and C domains. Why should any of the domains have higher stability? Perhaps because as a cell surface protein, SpA is subject to harsh conditions that favor unfolding and degradation and the more stable domains would be more resistant to these effects. A secreted multidomain virulence factor of *Streptococcus equisimilis*, streptokinase, also comprises domains that unfold independently with different stabilities. However, in that case, the least stable domain is actually the central domain rather than the N-terminal domain(58).

An alternative hypothesis is that the stability gradient plays a role in function. However, previous studies have shown that the five individual domains of SpA-N bind to Fc with similar affinities(11, 59). Another possibility is that the stability difference between domains helps avoid misfolding between domains(38). This is possible, but the N-to-C gradient is not necessary to achieve this effect. Clarke and coworkers have proposed that sequence identity would favor misfolding(60). However, as shown in Table 1, the sequence identity of the five domains in SpA-N is high. As described in Materials and Methods, the folding of SpA-N is highly reversible. Thus, it is unlikely that misfolding occurs.

The stability of E domain when connected to the rest of SpA-N corresponds to a longer linker between E and D domains (see Figure 1). There are several possible explanations for this enhanced stability. One consequence of the additional AQQ sequence could be that it nucleates helix and rigidifies the linker. However, when we compared the predicted helicity of the shorter sequence to the E– D linker sequence, the helicities were low and nearly identical. It is also unlikely that excluded volume effects in the unfolded state explain the stabilization of E, given that these effects would be at least as large in all the other domains as well. Although the E domain is 5 residues shorter than the others, the missing residues are not predicted to be helical and therefore are unlikely to provide stabilization. A hypothesis worth exploring is that E domain is stabilized because its linker is flexible and long enough to permit surface contacts between folded E and the rest of SpA-N. The lack of interaction between E and D observed by Starovasnik, et al.(33) may not apply when the E domain can interact with the entire SpA-N protein, which would provide a larger contact surface. This hypothesis could be tested with a modified SpA-N whose longer E–D linker is replaced by the conserved KADNKF linker.

In summary, we have measured the stabilities of the five antibody binding domains of SpA, both as isolated domains and in the context of full-length SpA-N. We have shown that four of the five domains, except E, have the same stability in both contexts, within error. This observation is consistent with the lack of structural(47) and thermodynamic coupling between the domains. Although the linkers between domains D through C are short (6 aa), our previous NMR and SAXS studies have demonstrated that they are highly flexible and therefore provide a thermodynamic buffer between domains. Because of the significant stability gradient from N-to C-terminus, the predicted thermal or chemical denaturation curves are broad, non-sigmoidal and relatively featureless.

## Materials and Methods

### Plasmid Construction

Oligonucleotides encoding WT SpA-N and the 5 fluorescent variants of SpA-N were synthesized using the method of Cox, et al.(61) The polyclonal DNA fragments were cloned into the pAED4 expression vector(62) after double digestion with NdeI and BamHI (New England Biolabs). These clones were then screened for expression of SpA-N by lysing IPTG induced cultures with BugBuster (Novagen) and adding IgG Agarose (Sigma) to the lysate. The mixture was incubated at room temperature with shaking for 30 minutes, and then centrifuged at 3000 x g for 2 minutes. The IgG Agarose pellet was washed 3 times with PBS, and the pellets were boiled in SDS-PAGE sample buffer and run on 4-20% TrisGlycine gels (Bio-Rad). Plasmids that overexpressed a 32 kDa protein were then submitted for DNA sequencing. EdpA, DdpA, AdpA, and CdpA were PCR cloned from the respective tryptophan substituted SpA-N variant. The PCR primers added a 5’ NdeI and a 3’ BamHI site, and were subsequently cloned into pAED4. BB, BBB, BBBB, and BBBBB were synthesized by GENEWIZ, Inc. These genes were synthesized in a pUC57 cloning vector, and were subsequently cloned into the pAED4 expression vector.

### Protein Expression and Purification

Plasmids were transformed into *Escherichia coli* BL21(DE3) cells using standard transformation procedure. A single colony of transformed bacteria was used to inoculate a 50 ml culture of LB media with 0.1 mg/ml ampicillin. This starter culture was incubated at 37°C until the O.D. reached 0.8-1.0, whereupon it was used to inoculate 1L cultures that were allowed to grow to O.D. 0.8-1.2 at 37°C. IPTG was then added to a final concentration of 1 mM and the cultures were incubated for an additional 6-8 hours. The cells were harvested by centrifugation and resuspended in 20-30 ml of 50 mM Tris pH 8.8, 1 mM EDTA, and protease inhibitor cocktail (AEBSF, pepstatin, bestatin and E-64). The cells were lysed in a French pressure cell and insoluble material was centrifuged from the lysate. The lysate was brought to pH 9.0 and micrococcal nuclease was added to digest large DNA fragments for 15 mins. The resulting solution was brought to 4M guanidinium HCl (BioBasic, Inc.) and 20 mM TCEP was added. In two successive steps, the solution was dialyzed in a 5% acetic acid buffer, which precipitated many cellular materials, but not expressed proteins. After centrifugation of the insoluble material the resulting solution was allowed to dialyze overnight into deionized water. The protein was further purified using two types of ion exchange chromatography. First, the protein was loaded onto a strong cation exchanging, SP Sepharose (GE) column in 50 mM acetate buffer at pH 3.6. The column was eluted with a NaCl gradient (typically 100 mM to 500 mM gradient) in a volume of 600-800 ml and collected in 10 ml fractions monitored by a UV detector (Bio-Rad) at 278 nm. The fractions comprising the protein elution peak were checked for purity by SDS-PAGE. The most pure fractions were pooled and dialyzed against deionized water. Subsequently, the resulting solution was loaded onto a weak anion exchanging DEAE Sephacel (GE) column in 25 mM Tris (BioBasic, Inc.) at pH 10.0 and eluted with a 800 ml NaCl gradient (typically 0 to 250 mM gradient) into 10 ml fractions monitored by a UV detector at 278 nm. The protein elution peak fractions were checked for purity by SDS-PAGE. The most pure fractions were pooled and dialyzed against deionized water. The final solution was lyophilized and stored in a desiccator. Purity of the final protein stocks was confirmed to exceed 95% by SDS-PAGE. The mass of all proteins was confirmed by ESI-MS.

### Fluorescence Detected Titrations

Lyophilized proteins were dissolved in deionized water to make stock solutions whose concentrations were calculated utilizing the method of Edelhoch(63). Urea (Nacalai Tesque) was used primarily as a denaturant because the folded and unfolded baselines were well sampled for most variants. In the cases where the unfolded baseline was not sampled well (C and B domains), we confirmed the extrapolated stabilities with that obtained using guanidinium chloride (GndCl), which is a stronger denaturant. The respective plot and corresponding thermodynamic parameters are in Supporting Information (Fig. S2 and Table S1). Urea (GndCl) titrations were done using a Hamilton automatic titrator and fluorescence spectra were collected with a Shimadzu RS-5301 spectrafluorimeter in 0.2M (0.1M) increments from 0M to 9.8M (6.0M) urea (GndCl) in a Perkin Elmer 1.0 cm fluorescence luminescence cuvette. Stock urea solutions were prepared fresh daily by dissolving in 20 mM HCl to minimize carbamylation of basic amino acid sidechains. The concentration of urea (GndCl) titrant was calculated using refractometry and the final 9.8M urea (6.0M GndCl) sample was also checked by refractometry. Final concentrations of 1 *µ*M protein were used for all urea titration measurements. Fluorescence spectra were collected using 50 mM sodium acetate, 100 mM NaCl pH 5.5 buffer. A 295 nm excitation wavelength was used to minimize the contribution from the tyrosine residues in neighboring domains to the fluorescence signals for the single tryptophan in a given domain in SpA-N. Identical conditions were used for single domains and multi-B domain variants for an equivalent comparison. Emission spectra were collected and summed from 345-350 nm, which is the low energy side of the tryptophan emission band, so that changes in the Stokes peak from the Raman effect of water did not obscure the results. Final titration curves are plotted as I/I0 normalized fluorescence. The thermodynamic analysis of all denaturation curves as well as the EdpA and E domain denaturation surface was done in Mathematica (Wolfram).

### Error Analysis

To estimate the reproducibility of the fitted parameters derived from the fluorescence-detected urea denaturation data the *m*_eq_ values for domains D, A, and B were compared between the two contexts for their differences. Because the domains are identical in both contexts, the *m*_eq_s should also be identical, but in fact they varied by 2.4%, 6.6%, and 4.3%. This leads to an estimated reproducibility standard deviation of 3.32%. Based on a Gaussian normal distribution, this gives 95% confidence intervals that are 1.96 times this value. This value was used to estimate the uncertainties of all parameters for all domains except that of E domain’s Δ*G*(0). The E domain Δ*G*(0) and *C*_1*/*2_ uncertainties come from the fits, with standard errors computed, allowing for correlated residuals.

### Circular Dichroism Spectroscopy

All far-UV CD spectra and temperature melts were collected using an Aviv 202S circular dichroism spectrapolarimeter (Aviv, Lakewood, NJ) equipped with a Peltier temperature controller. Far-UV CD spectra were collected in the range from 185-350 nm in a Hellma 0.1 mm quartz cell, with 200 *µ*M concentration for the single domain proteins. Concentrations were scaled according to the number of domains for multidomain proteins so that the concentration per domain remained 200 *µ*M. Spectra were collected at 25°C in a 50 mM sodium acetate, 100 mM NaCl pH 5.5 buffer, using a 1 nm bandwidth, 1 nm resolution, 3 second averaging time, and a 2 min pre-acquisition equilibration delay. Circular dichroism detected temperature melts were completed using a 1.0 cm four-sided polished fluorescence cuvette with an 8x3 mm stir bar used for proper stirring and temperature equilibration. We used a 5 *µ*M and 2.5 *µ*M protein concentration for B and BB, respectively, detecting the 222 nm CD signal for ten seconds at each temperature point from 2-97°C in 5°C increments with a temperature ramp speed of 10°C/min and a 2 min equilibration time. The buffer used for collecting CD spectra was used for these experiments. These repeats indeed overlaid with the 2 min equilibration time data. The denaturation surface data were collected by automated thermal melts in 5° steps in the presence of various urea concentrations ranging from 0 to 8M, depending on the stability of the domain. The urea concentrations were determined by refractive index.

### SpA-N denaturation

CD denaturation studies of SpA-N were carried out in the same Aviv CD instrument and in the same buffer described above. For the thermal melt, the wavelength was set at 222 nm with a 1 nm bandwidth. The temperature was ramped from 0 to 99°C in 1°C steps, with a 1-minute equilibration time between steps and 60-second signal averaging at each point. A reverse scan was executed to estimate the reversibility, which was 78.5%. This reversibility is typical for highly reversible proteins, given that at high temperature protein loss to the cell wall almost always occurs. Therefore, thermodynamic parameters derived from this denaturation curve were reliably associated with a reversible transition.

Urea denaturation was performed by preparing 14 individual samples in a urea range from 0 to 9.8 M. The CD signal at 222 nm was recorded at 25°C for 120 seconds at 1-second intervals, with a 0.5-second time constant, and the reported signal is the mean of that data. The urea concentrations of each sample were determined by refractive index.

The folded and unfolded baselines for both denaturation curves were fit to linear equations (see Figures S9 and S10). The thermodynamic parameters were held fixed at Tables 4 and 3 values, and only the baseline parameters were floated in the fits to the signal-based denaturation curves. For the urea denaturation analysis, the four baseline parameters were scaled by the per-domain signal weights. For the thermal denaturation analysis, the folded baseline parameters were floated, and the unfolded intercept was floated. However, the unfolded slope could not be left free because the unfolded state was not sufficiently populated. Instead, the unfolded baseline was reconstructed from the pre-transition folded line using the equation Σ_U_ = Σ_F_ + (Σ_obs_ − Σ_F_)*/f*_U_, which gave a corrected slope of −0.00734 *±* 0.00047 mdeg/°C that was held fixed.

### Use of Artificial Intelligence

Some of the data analysis was done with Claude Opus 5 (Anthropic), used through Claude Code. It was used to refit the fluorescence-detected urea denaturation curves. It wrote the code for fitting the SpA-N thermal and urea denaturation data. It detected a transposition of Δ*C*_*p*_ for domains E and D in Table 4. It wrote some of the code for generating Figures 9A, 9B, and S9 through S12. Claude assisted in the formatting of the manuscript, but all of the text was written by the authors. No AI-written prose is in this manuscript. The authors verified the calculations done by AI and are accountable for all quantitative and qualitative statements.

## Data, Materials, and Software Availability

Data and data analysis Wolfram notebooks are available in the Harvard Dataverse repository, https://doi.org/10.7910/DVN/J1XMN0.

## ACKNOWLEDGMENTS

The authors thank Jane Clarke, Yang Qi, Kan (Jonathan) Li and Roy Hughes for useful discussion. This work was supported by NIGMS grants R01-GM-081666 to TGO and R35-GM-144042 to BRD.

## Notes

https://doi.org/10.7910/DVN/J1XMN0

### Competing Interest Statement

The authors have declared no competing interest.

### Summary of Updates

Many changes to figures and tables. New experimental data for the thermal and chemical denaturation of SpA-N has been added. New error analysis for Tables 2, 3, and 4

## References

1. T.. Bae, et al., Staphylococcus aureus virulence genes identified by bursa aurealis mutagenesis and nematode killing. Proc. Natl. Acad. Sci. United States Am. 101, 12312 (2004).

2. JB.. Wardenburg, RJ.. Patel, O.. Schneewind, Surface proteins and exotoxins are required for the pathogenesis of staphylococcus aureus pneumonia. Infect. Immun. 75, 1040 (2007).

3. N.. Palmqvist, T.. Foster, A.. Tarkowski, E.. Josefsson, Protein a is a virulence factor in staphylococcus aureus arthritis and septic death. Microb. Pathog. 33, 239 (2002).

4. HK.. Kim, AG.. Cheng, HY.. Kim, DM.. Missiakas, O.. Schneewind, Isda and isdb antibodies protect mice against staphylococcus aureus abscess formation and lethal challenge. J. Exp. Medicine 207, 1863 (2010).

5. B.. Guss, et al., Region x, the cell-wall-attachment part of staphylococcal protein a. Eur. J. Biochem. 138, 413 (1984).

6. O.. Schneewind, A.. Fowler, KF.. Faull, Structure of the cell wall anchor of surface proteins in staphylococcus aureus. Science 268, 103 (1995).

7. O.. Schneewind, P.. Model, VA.. Fischetti, Sorting of protein a to the staphylococcal cell wall. Cell 70, 267 (1992).

8. T.. Moks, et al., Staphylococcal protein a consists of five igg-binding domains. Eur. J. Biochem. 156, 637 (1986).

9. S.. Lofdahl, B.. Guss, M.. Uhlen, L.. Philipson, M.. Lindberg, Gene for staphylococcal protein a. Proc. Natl. Acad. Sci. United States Am. Sci. 80, 697 (1983).

10. WW.. Navarre, O.. Schneewind, Proteolytic cleavage and cell wall anchoring at the lpxtg motif of surface proteins in gram-positive bacteria. Mol. Microbiol. 14, 115 (1994).

11. UK.. Ljungberg, et al., The interaction between different domains of staphylococcal protein a and human polyclonal igg, iga, igm and f(ab’)2: separation of affinity from specificity. Mol Immunol 30, 1279–85 (1993).

12. PK.. Peterson, J.. Verhoef, LD.. Sabath, PG.. Quie, Effect of protein a on staphylococcal opsonization. Infect. Immun. 15, 760 (1977).

13. S.. Sato, TL.. Religa, AR.. Fersht, Phi-analysis of the folding of the b domain of protein a using multiple optical probes. J. Mol. Biol. 360, 850 (2006).

14. SP.. Bottomley, et al., The stability and unfolding of an igg binding protein based upon the b domain of protein a from staphylococcus aureus probed by tryptophan substitution and fluorescence spectroscopy. Protein Eng. 7, 1463 (1994).

15. YW.. Bai, A.. Karimi, HJ.. Dyson, PE.. Wright, Absence of a stable intermediate on the folding pathway of protein a. Protein Sci. 6, 1449 (1997).

16. A.. Karimi, M.. Matsumura, PE.. Wright, HJ.. Dyson, Characterization of monomeric and dimeric b domain of staphylococcal protein a: Sources of stabilization of a 3-helix bundle protein. J. Pept. Res. 54, 344 (1999).

17. P.. Arora, TG.. Oas, JK.. Myers, Fast and faster: a designed variant of the b-domain of protein a folds in 3 µsec. Protein Sci. 13, 847 (2004).

18. G.. Dimitriadis, et al., Microsecond folding dynamics of the f13w g29a mutant of the b domain of staphylococcal protein a by laser-induced temperature jump. Proc. Natl. Acad. Sci. United States Am. 101, 3809 (2004).

19. P.. Arora, GG.. Hammes, TG.. Oas, Folding mechanism of a multiple independently-folding domain protein: Double b domain of protein a. Biochemistry 45, 12312 (2006).

20. S.. Sato, AR.. Fersht, Searching for multiple folding pathways of a nearly symmetrical protein: Temperature dependent ϕ-value analysis of the b domain of protein a. J. Mol. Biol. 372, 254 (2007).

21. J.. Deisenhofer, Crystallographic refinement and atomic models of a human fc fragment and its complex with fragment b of protein a from staphylococcus aureus at 2.9- and 2.8-a resolution. Biochemistry 20, 2361 (1981).

22. H.. Gouda, et al., Three-dimensional solution structure of the b domain of staphylococcal protein a: comparisons of the solution and crystal structures. Biochemistry 31, 9665 (1992).

23. M.. Tashiro, et al., High-resolution solution nmr structure of the z domain of staphylococcal protein a. J. Mol. Biol. 272, 573 (1997).

24. DY.. Zheng, JM.. Aramini, GT.. Montelione, Validation of helical tilt angles in the solution nmr structure of the z domain of staphylococcal protein a by combined analysis of residual dipolar coupling and noe data. Protein Sci. 13, 549 (2004).

25. M.. Graille, et al., Crystal structure of a staphylococcus aureus protein a domain complexed with the fab fragment of a human igm antibody: structural basis for recognition of b-cell receptors and superantigen activity. Proc. Natl. Acad. Sci. United States Am. 97, 5399 (2000).

26. LN.. Deis, et al., Multiscale conformational heterogeneity in staphylococcal protein a: possible determinant of functional plasticity. Structure 22, 1467–1477 (2014).

27. ZY.. Guo, CL.. Brooks, EM.. Boczko, Exploring the folding free energy surface of a three-helix bundle protein. Proc. Natl. Acad. Sci. United States Am. 94, 10161 (1997).

28. DOV.. Alonso, V.. Daggett, Staphylococcal protein a: unfolding pathways, unfolded states, and differences between the b and e domains. Proc. Natl. Acad. Sci. United States Am. 97, 133 (2000).

29. S.. Sato, TL.. Religa, V.. Daggett, AR.. Fersht, Testing protein-folding simulations by experiment: B domain of protein a. Proc. Natl. Acad. Sci. United States Am. 101, 6952 (2004).

30. PG.. Wolynes, Latest folding game results: protein a barely frustrates computationalists. Proc. Natl. Acad. Sci. United States Am. 101, 6837 (2004).

31. GG.. Maisuradze, A.. Liwo, S.. Oldziej, HA.. Scheraga, Evidence, from simulations, of a single state with residual native structure at the thermal denaturation midpoint of a small globular protein. J. Am. Chem. Soc. 132, 9444 (2010).

32. HX.. Lei, C.. Wu, ZX.. Wang, YQ.. Zhou, Y.. Duan, Folding processes of the b domain of protein a to the native state observed in all-atom ab initio folding simulations. J. Chem. Phys. 128, 235105 (2008).

33. MA.. Starovasnik, M. O’Connell, WJ.. Fairbrother, RF.. Kelley, Antibody variable region binding by staphylococcal protein a: Thermodynamic analysis and location of the fv binding site on e-domain. Protein Sci. 8, 1423 (1999).

34. MA.. Starovasnik, et al., Solution structure of the e-domain of staphylococcal protein a. Biochemistry 35, 15558–69 (1996).

35. S.. Batey, LG.. Randles, A.. Steward, J.. Clarke, Cooperative folding in a multi-domain protein. J. Mol. Biol. 349, 1045 (2005).

36. RS.. Gokhale, C.. Khosla, Role of linkers in communication between protein modules. Curr Opin Chem Biol 4, 22–7 (2000).

37. M.. Giorgi, CD.. Cianci, PG.. Gallagher, JS.. Morrow, Spectrin oligomerization is cooperatively coupled to membrane assembly: a linkage targeted by many hereditary hemolytic anemias? Exp Mol Pathol 70, 215–30 (2001).

38. KA.. Scott, A.. Steward, SB.. Fowler, J.. Clarke, Titin; a multidomain protein that behaves as the sum of its parts11edited by j. karn. J. Mol. Biol. 315, 819–829 (2002).

39. A.. Minajeva, M.. Kulke, JM.. Fernandez, WA.. Linke, Unfolding of titin domains explains the viscoelastic behavior of skeletal myofibrils. Biophys. J. 80, 1442 (2001).

40. LK.. Mosavi, S.. Williams, ZY.. Peng, Equilibrium folding and stability of myotrophin: A model ankyrin repeat protein. J. Mol. Biol. 320, 165 (2002).

41. T.. Aksel, D.. Barrick, Analysis of repeat-protein folding using nearest-neighbor statistical mechanical models in Methods in Enzymology. Vol. 455, p. 95 (2009).

42. ERG.. Main, Y.. Xiong, MJ.. Cocco, L D’Andrea, L.. Regan, Design of stable α-helical arrays from an idealized tpr motif. Structure 11, 497 (2003).

43. E.. Kloss, D.. Barrick, C-terminal deletion of leucine-rich repeats from yopm reveals a heterogeneous distribution of stability in a cooperatively folded protein. Protein Sci. 18, 1948 (2009).

44. DE.. Kamen, Y.. Griko, RW.. Woody, The stability, structural organization, and denaturation of pectate lyase c, a parallel β-helix protein. Biochemistry 39, 15932 (2000).

45. DW.. Bolen, MM.. Santoro, Unfolding free energy changes determined by the linear extrapolation method. 2. incorporation of delta g degrees n-u values in a thermodynamic cycle. Biochemistry 27, 8069 (1988).

46. JA.. Capp, A.. Hagarman, DC.. Richardson, TG.. Oas, The statistical conformation of a highly flexible protein: small-angle x-ray scattering of s. aureus protein a. Structure 22, 1184–1195 (2014).

47. Y.. Qi, et al., Continuous interdomain orientation distributions reveal components of binding thermodynamics. J. molecular biology 430, 3412–3426 (2018).

48. H.. Hjelm, K.. Hjelm, J Sjöquist, Protein a from staphylococcus aureus. its isolation by affinity chromatography and its use as an immunosorbent for isolation of immunoglobulins. FEBS letters 28, 73–76 (1972).

49. M.. Kubica, et al., A potential new pathway for staphylococcus aureus dissemination: the silent survival of s. aureus phagocytosed by human monocyte-derived macrophages. PLoS One 3, e1409 (2008).

50. PD.. Cotter, C.. Hill, Surviving the acid test: responses of gram-positive bacteria to low ph. Microbiol Mol Biol Rev 67, 429–53, table of contents (2003).

51. JK.. Myers, TG.. Oas, Preorganized secondary structure as an important determinant of fast protein folding. Nat Struct Biol 8, 552–8 (2001).

52. JH.. Han, S.. Batey, AA.. Nickson, SA.. Teichmann, J.. Clarke, The folding and evolution of multidomain proteins. Nat. Rev. Mol. Cell Biol. 8, 319–330 (2007).

53. GA.. Pellowe, et al., The human ribosome modulates multidomain protein biogenesis by delaying cotranslational domain docking. Nat. Struct. & Mol. Biol. 32, 2296–2307 (2025).

54. N.. Rajasekaran, CM.. Kaiser, Navigating the complexities of multi-domain protein folding. Curr. Opin. Struct. Biol. 86, 102790 (2024).

55. W.. Yu, D.. Missiakas, O.. Schneewind, Septal secretion of protein a in Staphylococcus aureus requires seca and lipoteichoic acid synthesis. eLife 7, e34092 (2018).

56. MA.. Catipovic, BW.. Bauer, JJ.. Loparo, TA.. Rapoport, Protein translocation by the seca atpase occurs by a power-stroke mechanism. The EMBO J. 38, EMBJ2018101140 (2019).

57. E.. Svetlicic, LA.. Alarcon, R.. Karlsson, C.. Jers, I.. Mijakovic, Mass spectrometry-based analysis of surface proteins in staphylococcus aureus clinical strains: Identification of promising k-mer targets for diagnostics. J. Proteome Res. 24, 4575–4585 (2025).

58. F Conejero-Lara, et al., Thermal stability of the three domains of streptokinase studied by circular dichroism and nuclear magnetic resonance. Protein Sci 5, 2583–91 (1996).

59. B.. Jansson, M Uhlén, PA.. Nygren, All individual domains of staphylococcal protein a show fab binding. FEMS Immunol Med Microbiol 20, 69–78 (1998).

60. CF.. Wright, SA.. Teichmann, J.. Clarke, CM.. Dobson, The importance of sequence diversity in the aggregation and evolution of proteins. Nature 438, 878 (2005).

61. JC.. Cox, J.. Lape, MA.. Sayed, HW.. Hellinga, Protein fabrication automation. Protein Sci. 16, 379 (2007).

62. DS.. Doering, “Functional and structural studies of a small F-actin binding domain,” PhD thesis, Massachusetts Institute of Technology (1992).

63. H.. Edelhoch, Spectroscopic determination of tryptophan and tyrosine in proteins. Biochemistry 6, 1948 (1967).

