## Supplemental Information for "An evolutionary strategy for highly efficient secretion of multidomain Staphylococcal protein A: N- to C-terminal stability gradient"

**Hagarman, *et al.***

The following supporting information includes a plot of the CD thermal denaturation of BdpA and BBdpA; a plot of the CD guanidinium chloride denaturation of BdpA and CdpA; a table reporting the $\Delta G_0$ and $m_{eq}$ estimates from fitting the GdnCl data; denaturation surface data and fits for all 5 isolated domains in 3D plot form; and plots depicting the baseline curves for SpA-N, thermal and chemical denaturations, and the corresponding microstate population plots for each.

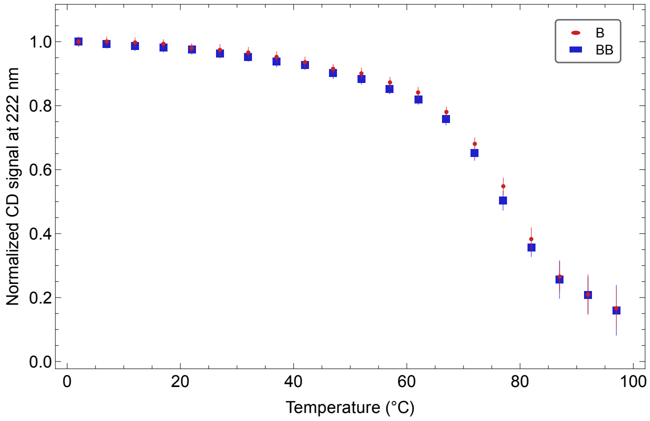


Figure S1: Circular dichroism detected temperature melts of single (red circles) and double (blue squares) B domain plotted in the temperature range 0–100 ℃. Error bars are included for each temperature point.

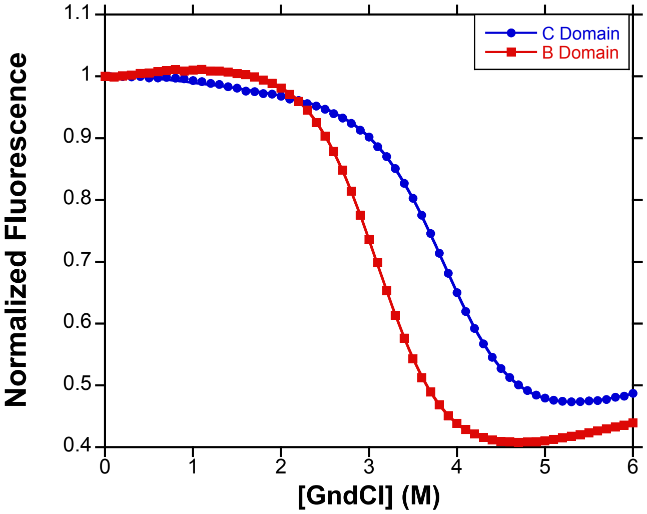


Figure S2: Fluorescence denaturation curves (points) and fits (solid lines) using GndCl as a denaturant for B (red squares) and C (blue circles) domains.

Table S1: The extrapolated stabilities and $m_{eq}$ values from B and C domain from guanidinium chloride denaturations. Uncertainties are estimated at 5% of the parameter values.

| Domain | BdpA | CdpA |
|---|---|---|
| $\Delta G(0)$ (kcal mol$^{-1}$) | $4.60 \pm 0.23$ | $5.6 \pm 0.28$ |
| $m_{eq}$ (kcal mol$^{-1}$ M$^{-1}$) | $-1.48 \pm 0.07$ | $-1.43 \pm 0.07$ |

### Denaturation Surface Analysis: The Definitions of $T_{hs}$ and $C_{hs}$

To obtain estimates for the key thermodynamic parameters of protein unfolding, it is necessary to perform both thermal and chemical denaturation in the same CD experiment. We have performed such experiments and have obtained estimates for the heat capacity, enthalpy, entropy and free energy of denaturation as well as the temperature-dependent $m_{eq}$. The data collected in this way form a three dimensional surface displaying CD signal as a function of temperature and urea concentration (*c.f.*, Fig.s S4-S8.) The CD denaturation surfaces of all 5 domains were fit to the following equations[**?**, **?**]:

$$\Delta G(T, [\text{Den}]) = \Delta H(T, [\text{Den}]) - T\Delta S(T, [\text{Den}]) \tag{1a}$$

$$\text{where: } \Delta H(T, [\text{Den}]) = \Delta C_p(T - T_{hs}) + h([\text{Den}] - C_{hs}) \tag{1b}$$

$$\text{and: } \Delta S(T, [\text{Den}]) = \Delta C_p \ln\left(\frac{T}{T_{hs}}\right) + s([\text{Den}] - C_{hs}) \tag{1c}$$

where $\Delta G, \Delta H,$ and $\Delta S$ are the free energy, enthalpy and entropy of unfolding and are all functions of temperature and urea concentration; $\Delta C_p$ is the heat capacity of unfolding; $h$ and $s$ are the enthalpic and entropic components of $m_{eq}$; $T_{hs}$ is the temperature of maximum stability; and $C_{hs}$ is the urea concentration such that $\Delta G(T_{hs}, C_{hs}) = 0$. Note that because $\Delta S$ is the temperature derivative of $\Delta G$, at $T_{hs}$ and $C_{hs}$, $\Delta S$ is also 0 and therefore, so is $\Delta H$. This is why, with $T_{hs}$ and $C_{hs}$ as reference conditions, Equations 1b and 1c have no reference values for $\Delta H$ and $\Delta S$. The $\Delta G(T, [\text{Den}])$ curves shown in Fig. S3 are derived from the parameter values fit to the AdpA denaturation surface (see Fig. S6) and demonstrate the utility of using $T_{hs}$ and $C_{hs}$ as reference conditions. It is at this temperature and [Den] where the change in signal is greatest and therefore carries the most information about the key thermodynamic parameters.

To fit the denaturation surfaces, it is necessary to use Equations 1 to compute the fraction unfolded and employ this along with equations for the folded and unfolded baseplanes:

$$F_{\text{unf}}(T, [\text{Den}]) = \frac{\exp\left(-\frac{\Delta G(T,[\text{Den}])}{RT}\right)}{1 + \exp\left(-\frac{\Delta G(T,[\text{Den}])}{RT}\right)} \tag{2a}$$

$$\Theta_{\text{F}} = \Theta_{\text{UF},0} + m_{\text{F,T}}T + m_{\text{F,Den}}[\text{Den}] \tag{2b}$$

$$\Theta_{\text{U}} = \Theta_{\text{U},0} + m_{\text{U,T}}T + m_{\text{U,Den}}[\text{Den}] \tag{2c}$$

$$\Theta(T, [\text{Den}]) = F_{\text{unf}}(T, [\text{Den}])\left(\Theta_{\text{U}} - \Theta_{\text{F}}\right) + \Theta_{\text{F}} \tag{2d}$$

where $F_{\text{unf}}$ is the fraction unfolded, $\Theta_{\text{F}}$ and $\Theta_{\text{U}}$ are the folded and unfolded baseplane equations with temperature slopes of $m_{\text{F,T}}$ and $m_{\text{U,T}}$, urea slopes of $m_{\text{F,Den}}$ and $m_{\text{U,Den}}$ and intercepts of $\Theta_{\text{F},0}$ and $\Theta_{\text{U},0}$, respectively. To identify good initialization values for the parameters in Equations 2b and 2c, data points that lie on each baseplane were identified by inspection and initial values for the baseplane parameters are obtained by fitting each to Equations 2b and 2c. With these as initial values, all data were fit to the thermodynamic parameters in Equations 1 and the baseplane parameters in Equations 2.

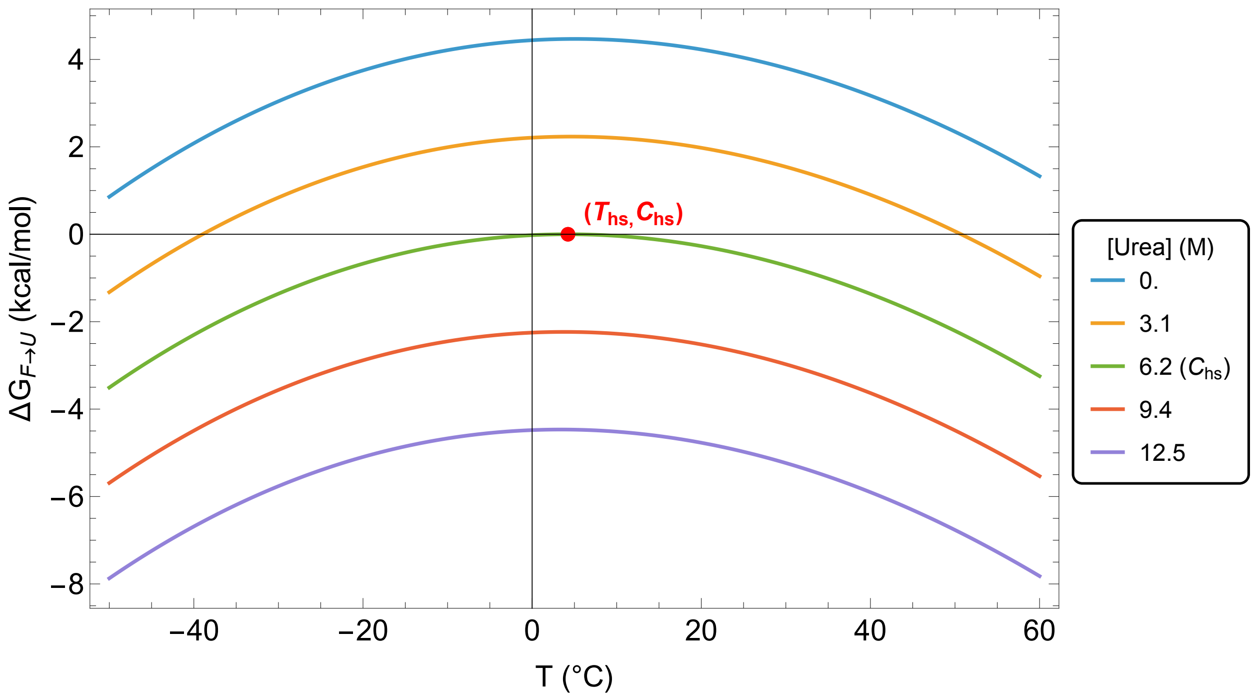


Figure S3: The free energy of unfolding as a function of temperature and urea concentration, using the values for AdpA listed in Table **??**. The red point marks the location of $\Delta G(T_{hs}, C_{hs})$.

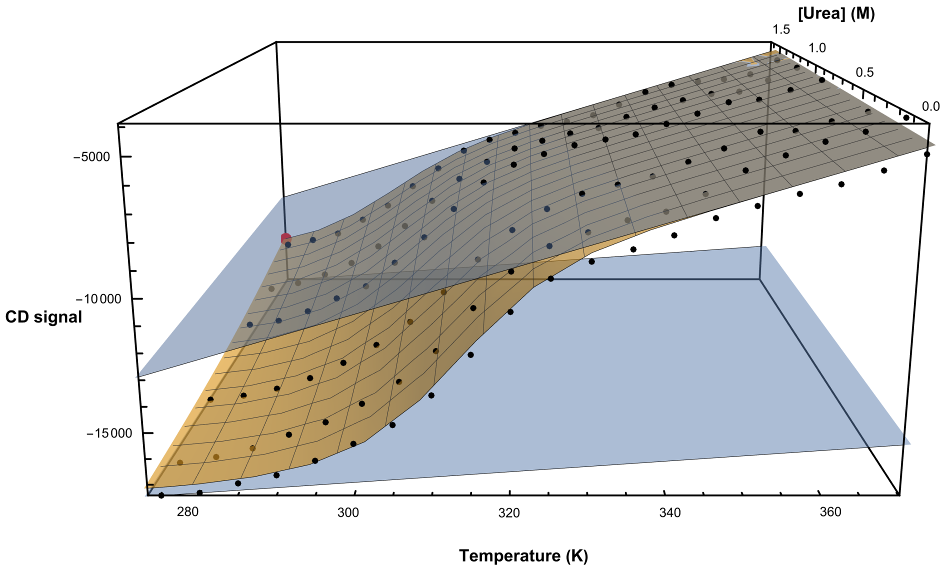


Figure S4: CD denaturation surface for EdpA fit to determine the parameter values listed in Table 4. The fitted denaturation surface is in gold, the fitted baseplanes are in blue. The observed CD signal are the black data points. The large red point corresponds to the predicted signal at $(T_{hs}, C_{hs})$, the point at which $\Delta G = \Delta H = \Delta S = 0$.

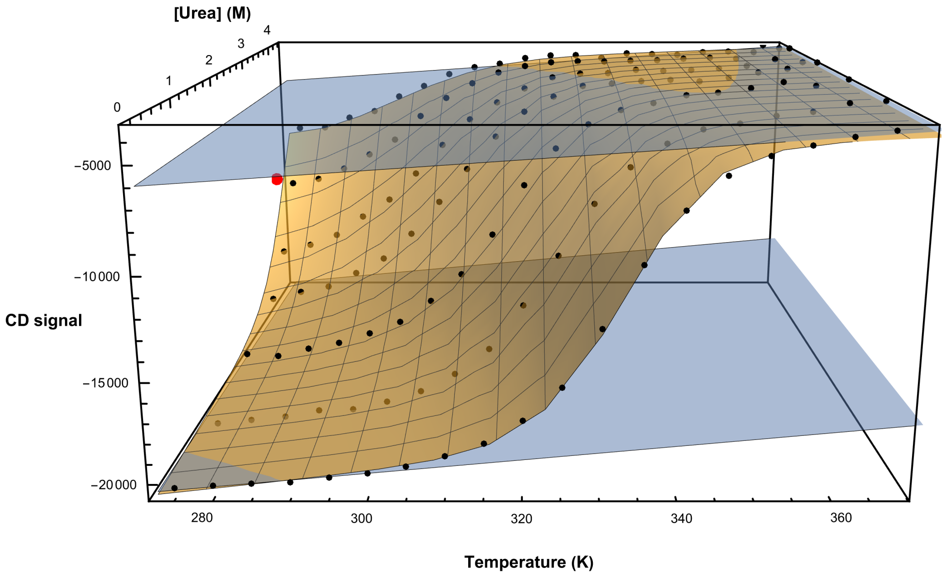


Figure S5: CD denaturation surface for DdpA fit to determine the parameter values listed in Table 4. The fitted denaturation surface is in gold, the fitted baseplanes are in blue. The observed CD signal are the black data points. The large red point corresponds to the predicted signal at $(T_{hs}, C_{hs})$, the point at which $\Delta G = \Delta H = \Delta S = 0$.

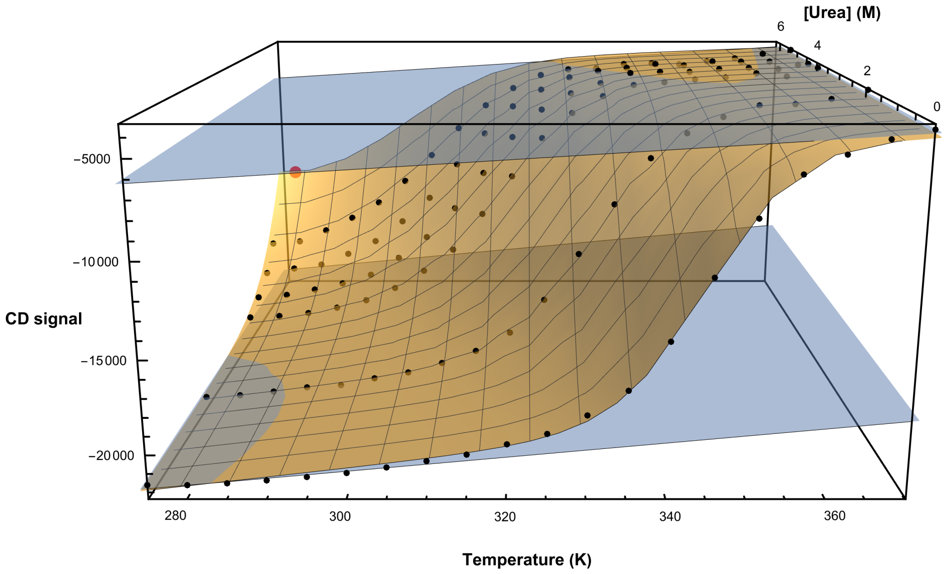


Figure S6: CD denaturation surface for AdpA fit to determine the parameter values listed in Table 4. The fitted denaturation surface is in gold, the fitted baseplanes are in blue. The observed CD signal are the black data points. The large red point corresponds to the predicted signal at $(T_{hs}, C_{hs})$, the point at which $\Delta G = \Delta H = \Delta S = 0$.

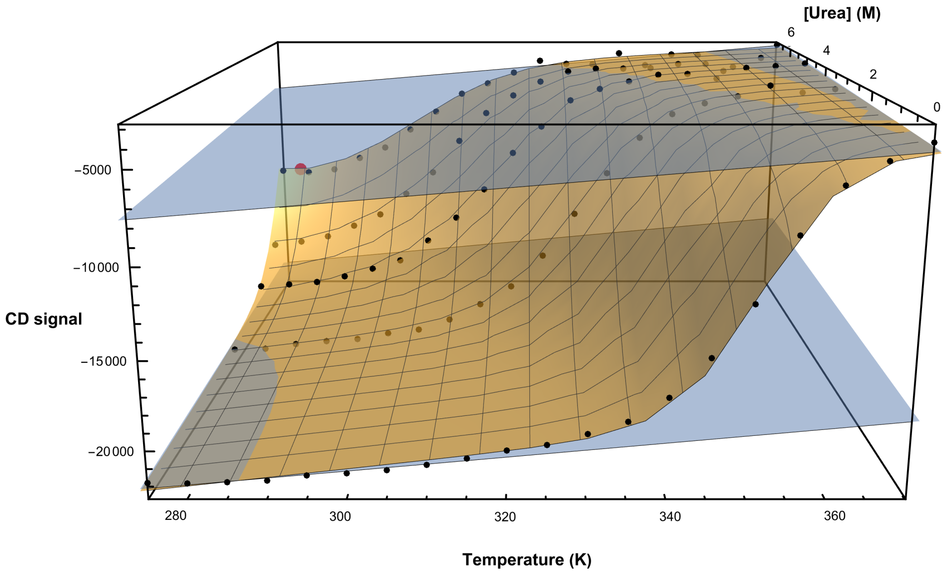


Figure S7: CD denaturation surface for BdpA fit to determine the parameter values listed in Table 4. The fitted denaturation surface is in gold, the fitted baseplanes are in blue. The observed CD signal are the black data points. The large red point corresponds to the predicted signal at $(T_{hs}, C_{hs})$, the point at which $\Delta G = \Delta H = \Delta S = 0$.

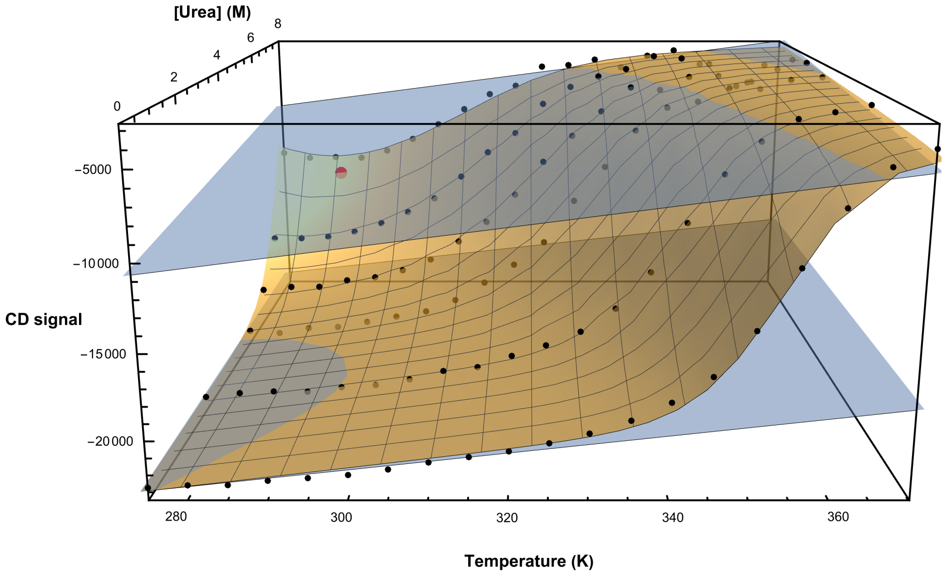


Figure S8: CD denaturation surface for CdpA fit to determine the parameter values listed in Table 4. The fitted denaturation surface is in gold, the fitted baseplanes are in blue. The observed CD signal are the black data points. The large red point corresponds to the predicted signal at $(T_{hs}, C_{hs})$, the point at which $\Delta G = \Delta H = \Delta S = 0$.

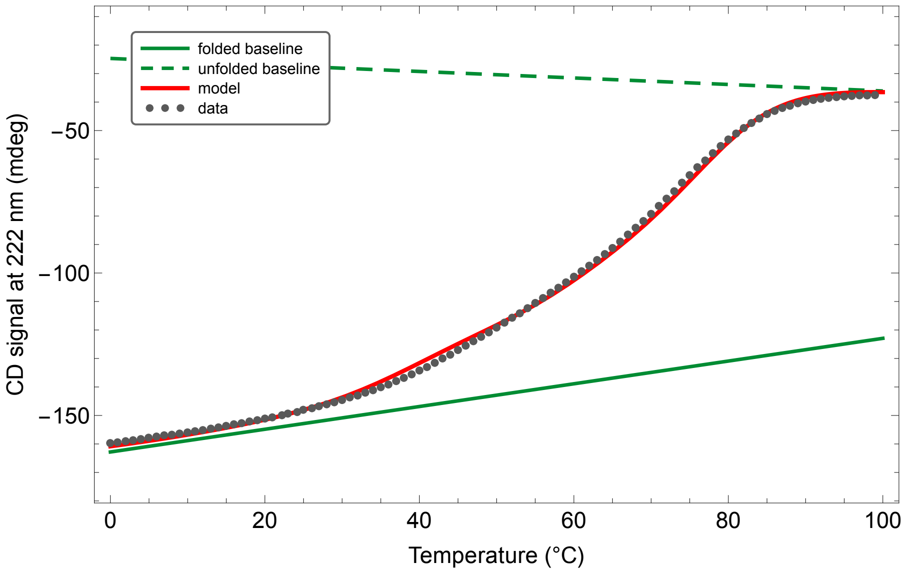


Figure S9: The CD signal of SpA-N versus temperature. The data were fit to the model whose parameters are listed in Table 4, combined with linear equations for the folded and unfolded baselines. The resulting baselines were used to convert the CD signal into fraction unfolded, as shown in Figure 9a.

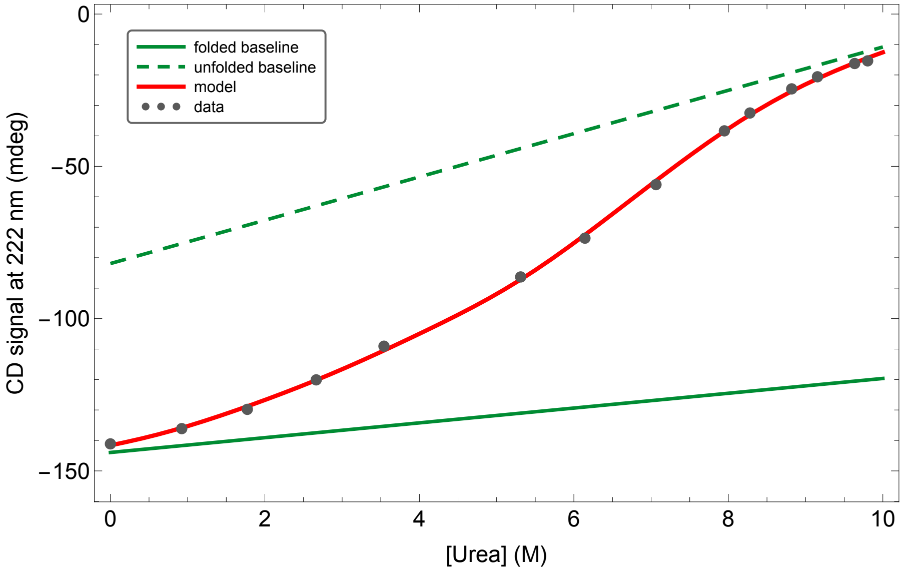


Figure S10: The CD signal of SpA-N versus urea concentration. The data were fit to the model whose parameters are listed in Table 3, combined with linear equations for the folded and unfolded baselines. The resulting baselines were used to convert the CD signal into fraction unfolded, as shown in Figure 9b.

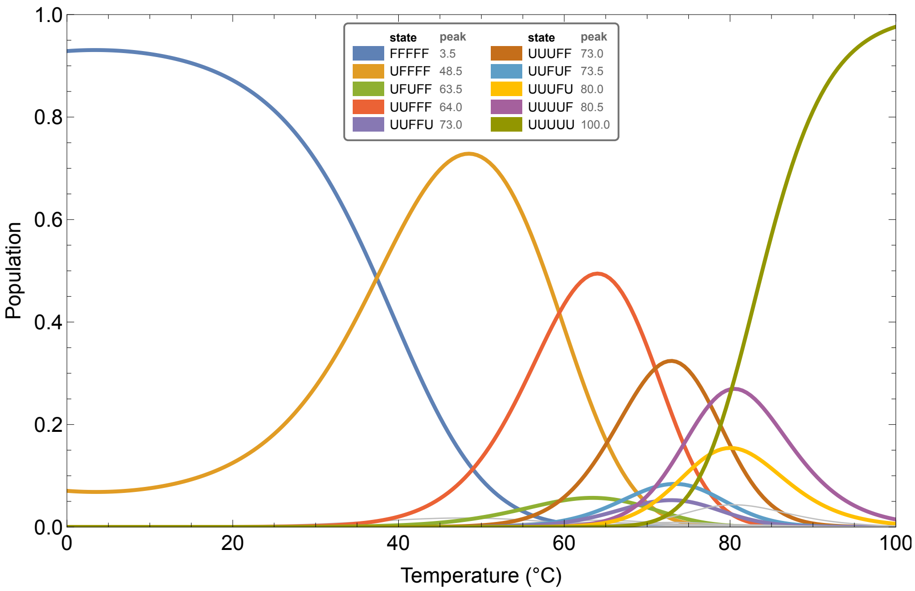


Figure S11: Populations of the microstates of SpA-N under thermal denaturation, based on the model whose parameters are listed in Table 4, the folded (F) and unfolded (U) states of each domain are indicated in the order EDABC

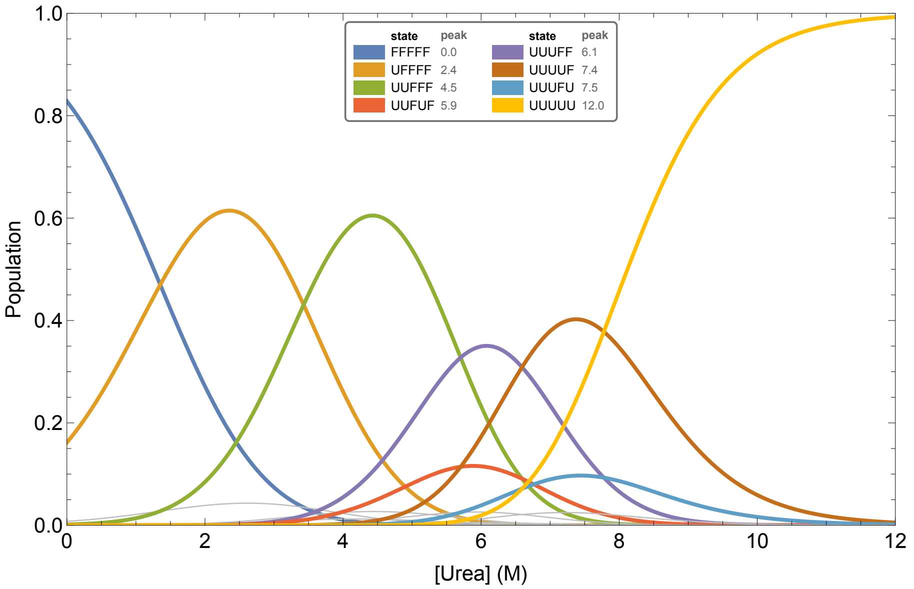


Figure S12: Populations of the microstates of SpA-N under urea denaturation, based on the model whose parameters are listed in Table 3, the folded (F) and unfolded (U) states of each domain are indicated in the order EDABC
